# Sense-and-response to intracellular signaling via nucleocytoplasmic localization control of a synthetic transcription factor platform

**DOI:** 10.64898/2026.10.07.757469

**Authors:** Pablo García-Díaz, Bhoomi Bhatt, Maria Teresa Morgado, Kayla Kim, Amara Okoye, Glenna Wink Foight

## Abstract

Many therapeutically relevant signaling pathways are sensed inside the cell, yet a modular synthetic biology platform for responding to intracellular signaling at the post-translational level is lacking. Signaling pathways control the nuclear localization and activity of native transcription factors via phosphorylation of nuclear localization/export or cytoplasmic anchor sequences. Here, we apply this concept to develop a platform of nucleocytoplasmic localization tags (NCtags) that control the nuclear translocation of a synthetic transcription factor (synTF) for transgene induction in engineered cells. We identify human-based synTF components that are basally cytoplasmic and induce transgenes under the control of NCtags with minimal background. We demonstrate the modular nature of this platform through transcriptional reporter induction in response to a calcium-inducible NFAT-derived NCtag, and development of novel NCtags responsive to hyperosmolarity, cAMP, or logical combinations of cAMP and calcium. Multiple factors in the tumor microenvironment elevate cAMP in T cells, leading to reduced function. We demonstrate that NCtag-synTF fusions expressed in T cells can respond to endogenous sources of cAMP or calcium in breast cancer spheroids to induce transgene expression. Our use of human components and an all-in-one lentiviral vector encoding the NCtag-synTF-transgene system establishes this as a novel platform for sense-and-response inside therapeutic cells.

## Introduction

Spatiotemporal control of transgene expression is an important strategy to improve the safety and efficacy of cell and gene therapy. Transgenes like cytokines, antibodies, chimeric antigen receptors (CARs), or metabolic factors may require spatial control to limit their expression to the site of disease and prevent toxic side effects. Dynamic control, i.e., dosage or temporal control, is also important to optimize efficacy, enabling precise levels of secreted factors or timing of transgene expression to coincide with a particular cell state or disease stage. Implementation of dynamic transgene expression commonly involves an inducible minimal promoter preceded by response elements bound by a native or synthetic transcription factor (synTF). Native TF-inducible systems, sometimes known as cell-state-responsive promoters, rely on endogenous signaling or transcriptional control of the TF to induce activity. This requires that the engineered cell type express that TF and limits transgene output levels to that dictated by the strength of the TF’s transcriptional activation domain (TAD). SynTFs, such as rTetR, contain DNA-binding domains (DBDs) chosen to be orthogonal to the host cell genome, meaning that their response elements are not present or are underrepresented. They must be constitutively expressed alongside the inducible transgene, and their induction comes from a post-translational controller. Choice of responsive controller, their orthogonal nature, and the ability to change transcriptional activation domain strength make synTFs highly modular for different applications. Despite these advantages, many synTFs have traditionally incorporated non-human components to achieve orthogonality, including the yeast Gal4 DBD and the viral VP64 TAD. The potential immunogenicity of these components may limit the persistence of engineered cells and hinder their therapeutic use.^1^ Optimization with human-derived DBDs and TADs could therefore improve clinical compatibility.^1–4^

Options to induce synTF activity include extrinsic or autonomously responsive controllers.^1,2^ Extrinsic controllers responsive to small-molecule drugs or light are useful for transgenes such as insulin^3,4^, glucagon-like peptide-1^5,6^, or cytokines like IL-12^7^, where there is a rationale for precise dosage and timing control by the patient, clinician, or a computer system. These systems usually act on a population; that is, they affect all cells with the synTF circuit in the same way and, in the case of small-molecule controllers, may have limited spatial control and slow on/off kinetics. Autonomous controllers put the induction timing in the hands of individual cells that intelligently sense a component of their environment or intracellular state to induce the synTF. Multiple receptor-based controllers induce synTF activity in response to extracellular soluble or membrane-bound ligands.^8^ These include the well-established synNotch system, and the humanized version, SNIPR, which have been used to sense tumor-associated antigens or soluble factors like VEGF and TGFβ, and induce CAR or cytokine transgenes to specifically direct CAR T cell activity against tumors.^9–12^ One limitation of many receptor-based controllers is an inability to program response to intracellular signals, which would be useful for sensing metabolites, kinase pathway activity, second messengers, and complex combinations indicative of therapeutic cell state.

The options for controllers that are autonomously responsive to intracellular inputs are limited. One class of autonomous synTF controller consists of post-translationally modified nucleocytoplasmic localization sequences. Often adapted from the disordered regulatory regions of native TFs, these controllers consist of nuclear localization or export sequences (NLS/NES) or binding sites for cytoplasmic anchor proteins like 14-3-3. The localization sequences are regulated by post-translational modifications (PTMs), primarily phosphorylation, which alters binding to importins, exportins, and anchor proteins, resulting in a shift of nucleocytoplasmic distribution. These regulatory region tags are genetically fused to the synTF, and signaling events that drive nuclear translocation of the tagged synTF induce transgene expression from the synTF-inducible promoter. One implementation of this concept used the regulatory region of murine NFAT2 (*NFATc1*) to control the nuclear localization of a Transcription Activator-Like Effector (TALE)-based synTF in HEK293T cells in response to calcium signaling.^13^ Fussenegger and colleagues have extended this concept in the Generalized Engineered Activation Regulators (GEARs) system, in which they employed regulatory regions from NFAT, NF-κB (p65), SMAD2, or Elk-1 to control a dCas9-based activation system, inducing insulin and IL-12 expression in cell lines in response to pathway activation by exogenously sourced extracellular ligands.^13^ These proof-of-concept studies illustrated the potential utility of localization-based controllers, but their relevance to cell therapy is limited by their use of large, immunogenic synTFs and failure to implement their systems in primary cells with clinically relevant vectors. Additionally, they do not define the properties of synTFs that allow them to be controlled on the level of nucleocytoplasmic localization or determine how to engineer regulatory regions more generally.

Here, we extend the concept of localization-based controllers into a modular platform of synTFs and regulators. We call this class of controllers nucleocytoplasmic localization tags (NCtags) and profile the response of multiple NCtags to T-cell stimulation. We identify a synTF composed of a human DBD and TAD that can be controlled by NCtags, and define the properties that determine synTF controllability, establishing design principles for localization-controlled synTFs. Demonstrating the extensibility of the NCtag concept, we identify novel NCtags that respond to different combinations of calcium- or cyclic-AMP (cAMP)-signaling, or hyperosmolarity, expanding the range of intracellular inputs that can be converted into transgene output. For translation, we implement the platform in primary human T cells, encode the system in clinically relevant gamma retroviral and lentiviral vectors, and demonstrate CAR T cells that sense and respond to endogenous signaling sources in the tumor microenvironment. The NCtag and synTF platform utilizes a modular and extensible architecture and provides a novel post-translational control strategy for sense-and-response to intracellular signaling states.

## Results

### NCtags translocate in response to signaling downstream of TCR stimulation

We set out to build a post-translational control platform of NCtags that sense intracellular signaling states and respond by controlling the nuclear localization and activity of a synTF to drive transgene expression (Fig. 1A). We worked directly in primary human T cells wherever possible, restricting Jurkat T cells to preliminary screening of large construct panels, as signaling is often aberrant in immortalized cell lines. To rapidly profile NCtag and synTF localization, we developed a workflow for assessing mGreenLantern-tagged (hereafter “GFP”) constructs in fixed T cells by confocal microscopy in 96-well plates (Supplementary Fig. 1A).^14^ This requires fewer cells and quantifies samples faster than flow-based imaging cytometry. As T cells are suspension cells with low cytoplasmic volume, we compared quantification metrics and found that the correlation of GFP with Hoechst nuclear stain produced a larger effect size with lower variation than the cytoplasmic/nuclear GFP ratio commonly used for adherent cells (Supplementary Fig. 1B-D).

**Fig. 1.**
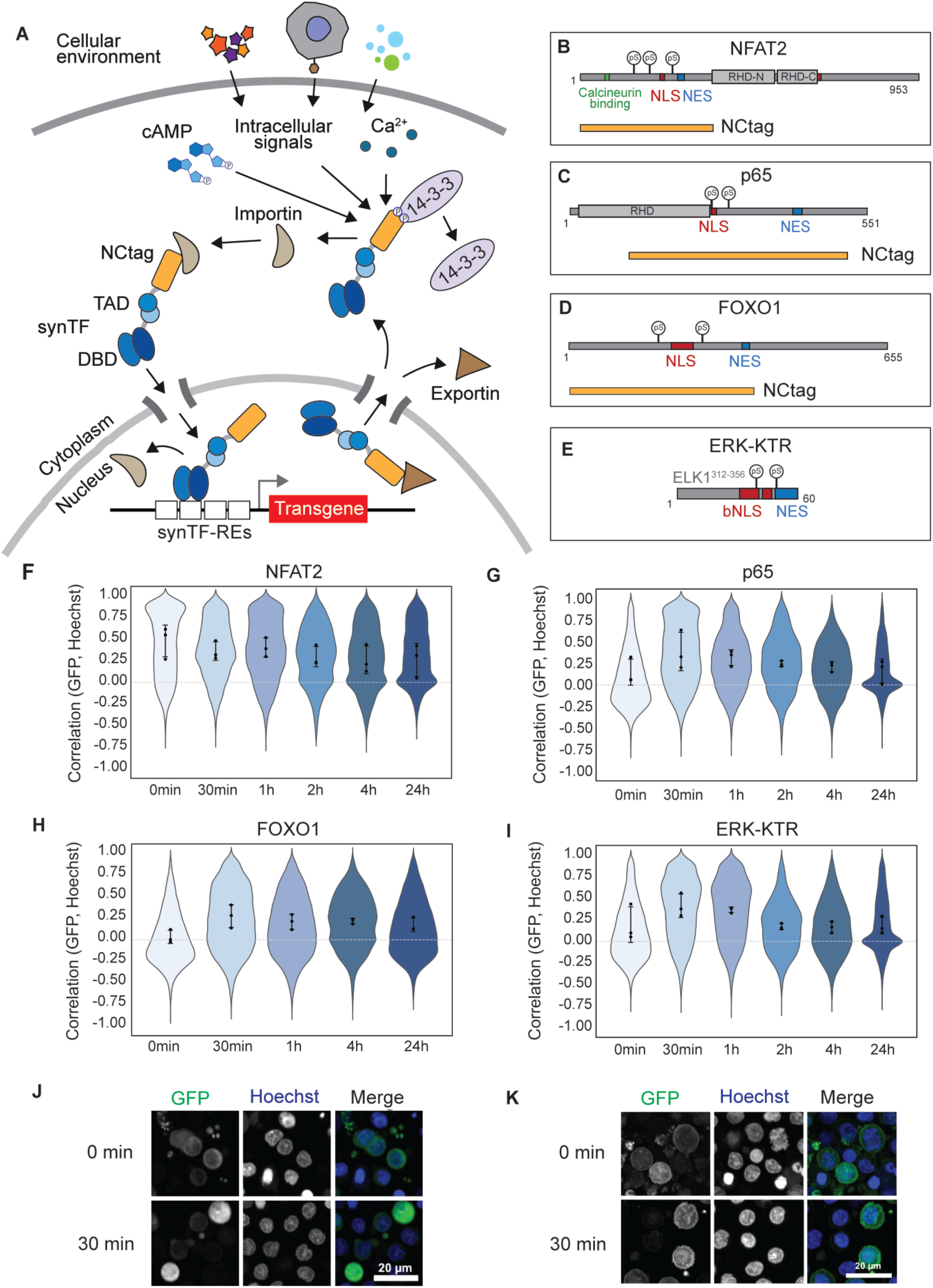
NCtags exhibit signaling-responsive localization change in T cells. **A** Schematic of the NCtag-synTF platform. Extracellular ligands generate intracellular signals that regulate the nuclear-cytoplasmic localization of an NCtag–synTF fusion. Domain schematics of NCtags derived from **B** full-length NFAT2, **C** full-length p65 (RHD, Rel-Homology domain), **D** full-length FOXO1, and **E** ERK-KTR based on ELK1. Orange bar indicates region used as the NCtag (all for ERK-KTR). pS, phosphoserine. Violin plots of single-cell GFP-Hoechst correlation quantified from confocal microscopy images for **F** NFAT2, **G** p65, **H** FOXO1, and **I** the ERK-KTR NCtags in primary T cells stimulated with plate-bound anti-CD3/CD28 and imaged at indicated times. Violins represent >9,000 cells per condition; dots indicate per-donor means (n=3 donors); error bars show SD. Representative images of **J** FOXO1 and **K** ERK-KTR NCtags in primary T cells under no-treatment and 30 min after anti-CD3/CD28 stimulation.

We used this workflow to profile a panel of putative NCtags for localization changes induced by TCR/CD28 stimulation with plate-bound anti-CD3/anti-CD28 antibodies in primary T cells. T cell stimulation activates a broad range of pathways, providing a general stimulus likely to translocate NCtags responsive to many kinases.^15^ Our panel included NCtags derived from human FOXO1, NFAT2, and p65, as well as an engineered kinase translocation reporter (KTR) that responds to ERK signaling (Fig. 1B-E).^13,16–18^ Quantification of NCtag-GFP localization over 24 h of stimulation showed that the NFAT2-NCtag was highly variable, with a subpopulation gradually shifting out of the nucleus (Fig. 1F), while FOXO1-NCtag, p65-NCtag, and ERK-KTR showed rapid nuclear translocation (high GFP-Hoechst correlation) followed by gradual return to the cytoplasm (low correlation) (Fig. 1G-K). The results for ERK-KTR and NFAT2 were unexpected, given that ERK-KTR is reported to translocate from the nucleus to the cytoplasm in response to ERK signaling and NFAT translocates from the cytoplasm to the nucleus in response to calcium signaling.^16,21,22^ The observed localization may reflect differences in basal signaling activity in the primary T cells and the complexity of signaling activated by combined TCR and CD28 activation. These preliminary tests demonstrated that multiple native and engineered post-translational regulatory regions translocate from the cytoplasm to the nucleus in response to signaling in T cells, demonstrating the potential to induce nuclear translocation of a fused synTF.

### Development of a human-based synTF for localization control

We designed a panel of human DBDs and TADs to identify synTF components without nuclear localization bias, evaluating their localization and control of an mScarlet transgene in Jurkat cells. Hypothesizing that the DBD would have a stronger effect on localization than the TAD, we first tested a panel of DBDs fused to the VP64 TAD and GFP in four constitutive localization variants: no tag, three NLS copies, three NES copies, or a CAAX membrane-targeting motif. We generated reporter constructs for each DBD by placing the DBD-specific response elements (REs) upstream of a synthetic minimal promoter (SMP) driving mScarlet expression.^19,20^ DBD-GFP-VP64 and reporter constructs were cotransduced into Jurkat cells using gamma retrovirus.

To maximize mScarlet dynamic range, we looked for DBDs that showed the highest output in the NLS control and lowest output in the NES control. Homeodomain DBDs derived from the liver-specific hepatocyte nuclear factor 1 homeobox A (HNF1A)^10,21^, the paired box 6 Pax6 (PAX6) proteins^10^, de novo-designed DBP35^22^, and a 3-finger zinc finger, ZF3^23^, showed higher mScarlet expression in the NLS control compared to the NES control, as desired. The GAL4 DBD also displayed this pattern, though the GFP-synTF expression levels were low compared to other DBDs (Supplementary Fig. 2A). Conversely, ZFHD1, a fusion of zinc fingers 1 and 2 from Zif268 and a homeodomain from Oct-1^24^, and the 6-finger zinc fingers ZiFTR3 and ZiFTR10^25^ showed higher mScarlet expression in the NES control (Fig. 2A). The construct expression levels may explain some of the mScarlet output patterns, as illustrated by the HNF1A-DBD No Tag control, where both GFP and mScarlet expression were higher than in the NLS control (Fig. 2B, Supplementary Fig. 2A), but we suspected that synTF localization was the strongest determinant.

**Fig. 2.**
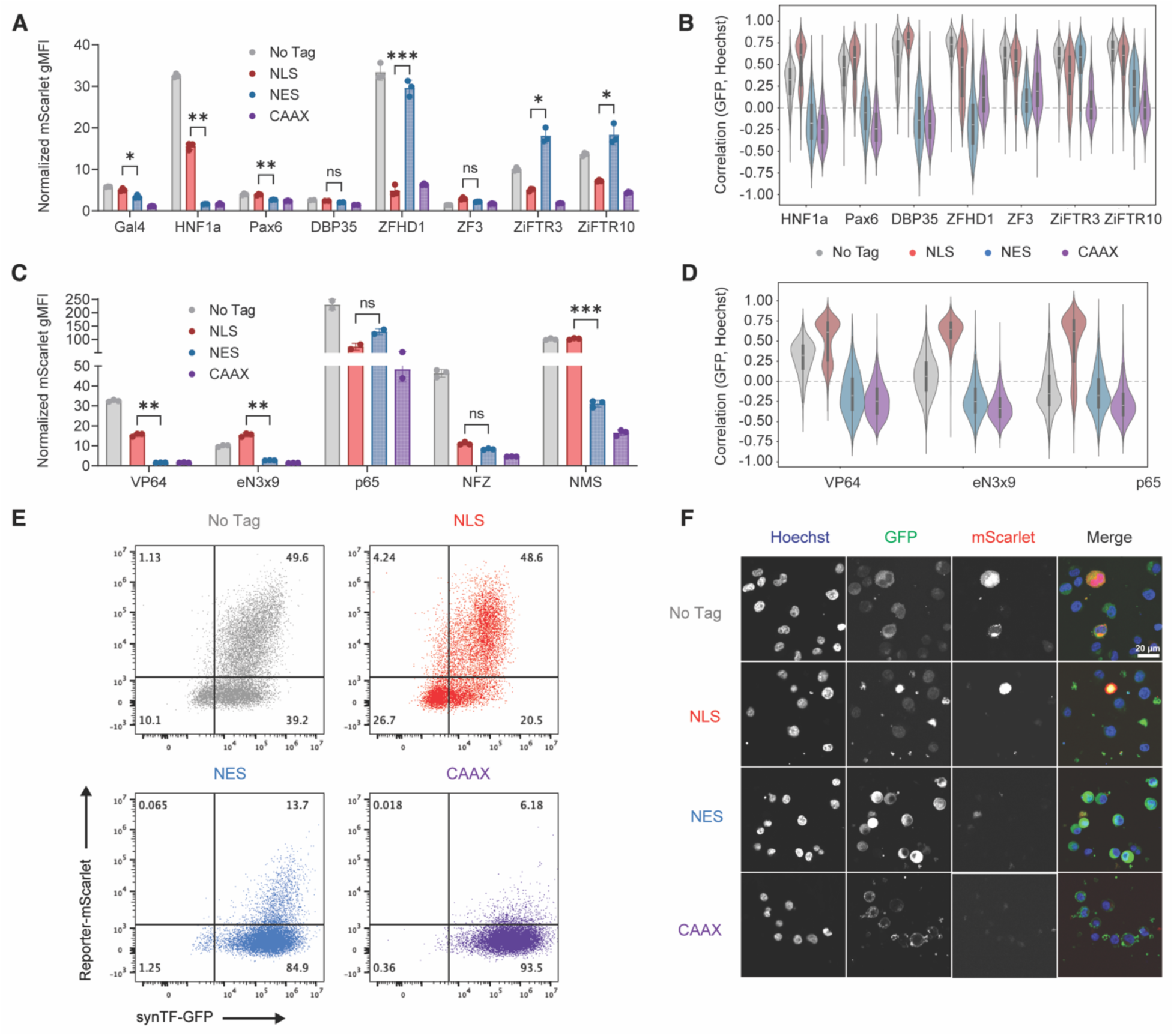
Development of a human-based synTF for localization control. All panels show Jurkat cells. **A** Reporter mScarlet geometric mean fluorescence intensity (gMFI) output of GFP⁺ cells expressing DBDs fused to VP64 and the indicated localization controls. **B** Violin plots of single-cell GFP/Hoechst correlation quantified from confocal microscopy images of DBD-VP64 constructs with the indicated localization controls. **C** Reporter mScarlet gMFI output of GFP⁺ cells expressing TADs fused to HNF1A-DBD and the indicated localization controls. **(A, C)** mScarlet gMFI normalized to untransduced; n = 3 separate transductions (n = 2 for p65); error bars show SD; significance calculated by two-way ANOVA with Tukey’s adjustment for multiple comparisons; * p < 0.05, ** p < 0.01, *** p < 0.001. **D** Violin plots of single-cell GFP/Hoechst correlation quantified from confocal microscopy images of TADs fused to HNF1A-DBD with the indicated localization controls. **(B, D)** Violins represent >140 cells per construct from four technical replicates of a single transduction; horizontal line indicates median; black bar indicates 25th–75th percentiles; whiskers indicate 1.5× the interquartile range. **E** Representative flow cytometry dot plots depicting GFP construct expression and mScarlet reporter output of HEsynTF fused to the indicated localization controls from a single transduction. **F** Representative confocal microscopy images of the constructs in E.

Using our confocal microscopy workflow, we quantified the subcellular localization of all DBD candidates (Fig. 2B). HNF1A, PAX6, and DBP35 displayed the expected localization pattern across all controls, matching their mScarlet output profiles; Gal4 expression was too low to quantify localization. However, the zinc-finger DBDs displayed varying degrees of mislocalization, particularly the NES constructs. For ZFHD1, the NES control showed a GFP-Hoechst correlation below 0, indicating cytoplasmic localization, yet retained high mScarlet expression, potentially due to high GFP-synTF expression (Supplementary Fig. 2A). ZiFTR3 and ZiFTR10 showed nuclear localization in the NES control (correlation values above 0), with ZiFTR3-NES exhibiting even higher nuclear localization than the NLS control, consistent with their mScarlet expression patterns. ZF3 showed weak nuclear localization in the NES control, though the NLS control retained higher nuclear localization, consistent with its mScarlet profile.

Overall, the homeodomain DBDs derived from HNF1A and PAX6 displayed a broader dynamic range in nucleocytoplasmic localization and transcriptional output compared to zinc-finger DBDs. Seeking a mechanistic basis for this, we found that DBD isoelectric point (pI) correlated with the differential localization between NLS and NES constructs and with the basal localization of the No Tag constructs (Supplementary Fig. 2B). Homeodomains and DBP35 (low pI) exhibited a larger dynamic range of localization than zinc fingers (high pI). We hypothesize that zinc fingers, which are positively charged at neutral pH, may exhibit nonspecific affinity for negatively charged DNA, driving basal nuclear localization, and potentially making them difficult to localize to the cytoplasm. Based on its high localization dynamic range and strong transcriptional output, we selected HNF1A as the DBD for our synTF.

We next evaluated human TADs paired with the HNF1A-DBD with the localization controls (Fig. 2C). The TADs of p65 and NMS (a chimeric fusion of TADs from NRF2, MRTF-A, and STAT1)^26^ showed high mScarlet output in the NLS control. However, p65 showed high background, with the NES control exhibiting high mScarlet. NMS yielded high mScarlet in the NES and CAAX controls, but gave the expected localization-dependent pattern. In contrast, eN3x9 (a small fusion of TADs from NRF2, MRTF-B, and MYOCD)^26^ showed mScarlet output in the NLS control comparable to VP64; while NFZ (a fusion of TADs from NCOA3, FOXO3, and ZN473)^27^ exhibited non-significant difference between the NLS and NES control and low GFP expression compared to other TADs (Supplementary Fig. 2C). The No Tag control from eN3x9 showed lower mScarlet output than VP64 despite similar GFP construct expression levels. Consistent with reporter output, the eN3x9 TAD showed less nuclear localization than VP64 for the No Tag control (Fig. 2D). p65 TAD showed the expected localization profile, with even more cytoplasmic localization for the No Tag construct than en3x9; the extremely high expression levels of the p65 constructs may explain the lack of localization-dependent mScarlet. All TADs had low pI values, suggesting that TAD pI is not a strong determining factor of localization (Supplementary Fig. 2D).

We chose HNF1A-DBD paired with eN3x9 (hereafter “HEsynTF”) as an optimal synTF for nucleocytoplasmic control with NCtags as it showed the expected pattern of reporter output (Fig. 2E) and localization (Fig. 2F) with minimal reporter expression when localized to the cytoplasm. With careful expression tuning, HNF1A-p65 may also be a suitable synTF for NCtag control. Due to the sourcing of DBD and TAD from human proteins, HEsynTF is predicted to contain few non-human, immunogenic peptides that could be displayed on a class I MHC, substantially less than predicted for a CAR (Supplementary Fig. 2E).

### Control of the synTF by calcium-responsive NCtags

We next evaluated whether HEsynTF could be controlled by calcium-responsive NCtags. To do this, we fused the regulatory regions (rr) of the four calcium-responsive NFAT-family members (human NFAT1–4) to the HEsynTF (Fig. 3A). Primary human T cells were first transduced with the HEsynTF retroviral reporter and subsequently with the HEsynTF-NFAT1–4rr panel. After stimulation with plate-bound anti-CD3 and anti-CD28 antibodies to activate TCR and CD28 signaling, we quantified localization and observed that NFAT1rr and NFAT3rr showed an increase in correlation after 1 h but rapidly returned to baseline (Fig. 3B,C). NFAT2rr and NFAT4rr showed a progressive decrease in correlation (Fig. 3B,D), consistent with our earlier results with NFAT2rr-GFP (Fig. 1F). Despite the transient nature of nuclear localization, all four NFAT regulatory regions induced reporter output (Fig. 3E,F). Given the single-cell variability and timing dependence of the localization readout, we therefore relied primarily on reporter output as a more robust indicator of NCtag-synTF functionality for further NCtag development.

**Fig. 3.**
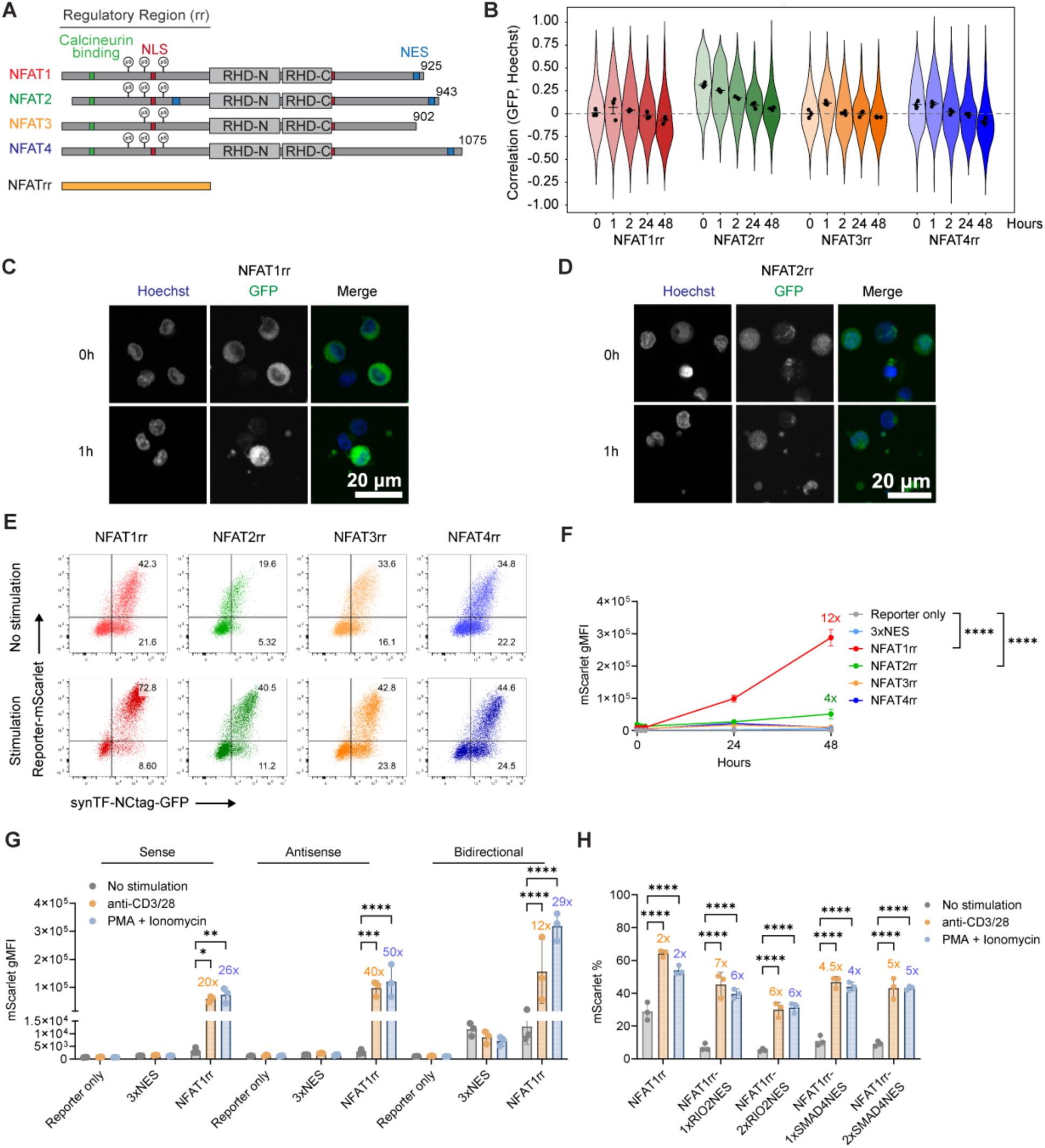
Control of synTF by calcium-responsive NCtags. All panels show primary T cells expressing HEsynTF fused to the indicated NCtags, unless otherwise noted. **A** Domain **s**chematic of NCtags derived from the regulatory regions (rr) of the calcium-responsive NFAT transcription factors. Orange bar indicates region used as the NCtag. **B** Violin plots of single-cell GFP-Hoechst correlation quantified from confocal microscopy images of primary T cells expressing NFAT-NCtags stimulated with plate-bound anti-CD3/CD28 and imaged at indicated times. Violins represent >135 cells per condition; dots indicate per-donor means (n=3 donors); error bars show SD. Representative images of primary T cells expressing, **C** NFAT1rr-synTF, or **D** NFAT2rr-synTF after 0 hr or 1 hr of anti-CD3/CD28 stimulation (mScarlet not shown). **E** Representative flow cytometry dot plots of cells showing the GFP expression and mScarlet reporter output upon stimulation with anti-CD3/CD28 for 48 h of one donor. **F** Reporter mScarlet gMFI output of GFP⁺ cells stimulated by anti-CD3/CD28 for the indicated time. n = 3 donors, mean ± SD; significance calculated by two-way ANOVA with Tukey’s adjustment for multiple comparisons. **G** Reporter mScarlet gMFI output of GFP⁺ cells expressing the indicated constructs with different lentiviral architectures after 24 h stimulation with the indicated treatments. **H** Percent mScarlet⁺ reporter output in GFP⁺ cells expressing HEsynTF-NFAT1rr fused to different NES sequences (with WPRE removed from the vector) after 24 h stimulation with the indicated treatments. (**G,H**) n = 3 donors, error bars show SD; significance calculated by two-way ANOVA with Dunnett’s adjustment for comparisons vs. no stimulation (for all panels unlabeled comparisons are non-significant and, * p < 0.05, ** p < 0.01, *** p < 0.001, **** p < 0.0001).

The NFAT1rr NCtag produced the strongest induction after 48h, yielding a 12-fold increase in reporter activation relative to 0h. NFAT2rr generated the next highest response at 4-fold activation. As GFP construct expression also increased with T cell stimulation, consistent with the global upregulation of RNA and protein synthesis that accompanies lymphocyte activation^28^, we used the 3xNES control to account for this effect; despite showing the highest GFP expression increase with stimulation, the 3xNES control exhibited low mScarlet, confirming that reporter induction is NCtag-dependent (Supplementary Fig. 3A).

To enable delivery with other vectors such as a CAR, we combined the NCtag-synTF and reporter into an all-in-one lentiviral vector, testing sense, antisense, and bidirectional configurations for NFAT1rr, 3xNES, and reporter-only controls (Supplementary Fig. 3B). We performed 24-hour anti-CD3/CD28 stimulation or PMA/IO treatment to induce NFAT1rr activation in primary T cells. Although the antisense vectors generated the highest fold-change in reporter expression (Fig. 3G), they also exhibited elevated basal activity compared to the sense vectors (Supplementary Fig. 3C). The bidirectional configuration showed higher basal activity and lower fold-change in reporter expression compared to the sense vectors. Based on this, we selected the sense configuration as the optimal vector for subsequent experiments (Fig. 3G, Supplementary Fig. 3C).

To minimize background induction, we added one or two copies of human NES sequences to the NFAT1rr vector, an NES with high nuclear export activity from RIO2 or with moderate activity from SMAD4.^29^ After 24-hour stimulation, these modifications lowered basal activity, reducing both mScarlet-positive percentage and gMFI (Fig. 3H, Supplementary Fig. 3D,E), and increased induction fold change compared to NFAT1rr alone. The RIO2 NES showed a lower background than the SMAD4 NES, consistent with its stronger nuclear export activity, and 1×RIO2-NES achieved the highest fold change in both mScarlet percentage and gMFI.

### Development of a hyperosmolarity-sensing NCtag

We next examined whether HEsynTF could be controlled by NCtags derived from NFAT5, which is reported to respond to hyperosmolarity by nuclear translocation and condensation.^30,31^ We designed a panel of NCtags (Fig. 4A) fused only to the HNF1A-DBD, as the NCtags contain the endogenous TADs of NFAT5 required for hyperosmolarity response.^30,31^ We first tested the panel in Jurkat cells with culture medium supplemented to a final concentration of 190 mM NaCl (chosen based on reported NFAT5 responsiveness)^32,34^ and observed that v9 and v11 yielded the highest reporter fold change (Fig. 4B) and showed an increase in nuclear localization (Fig. 4C).

**Fig. 4.**
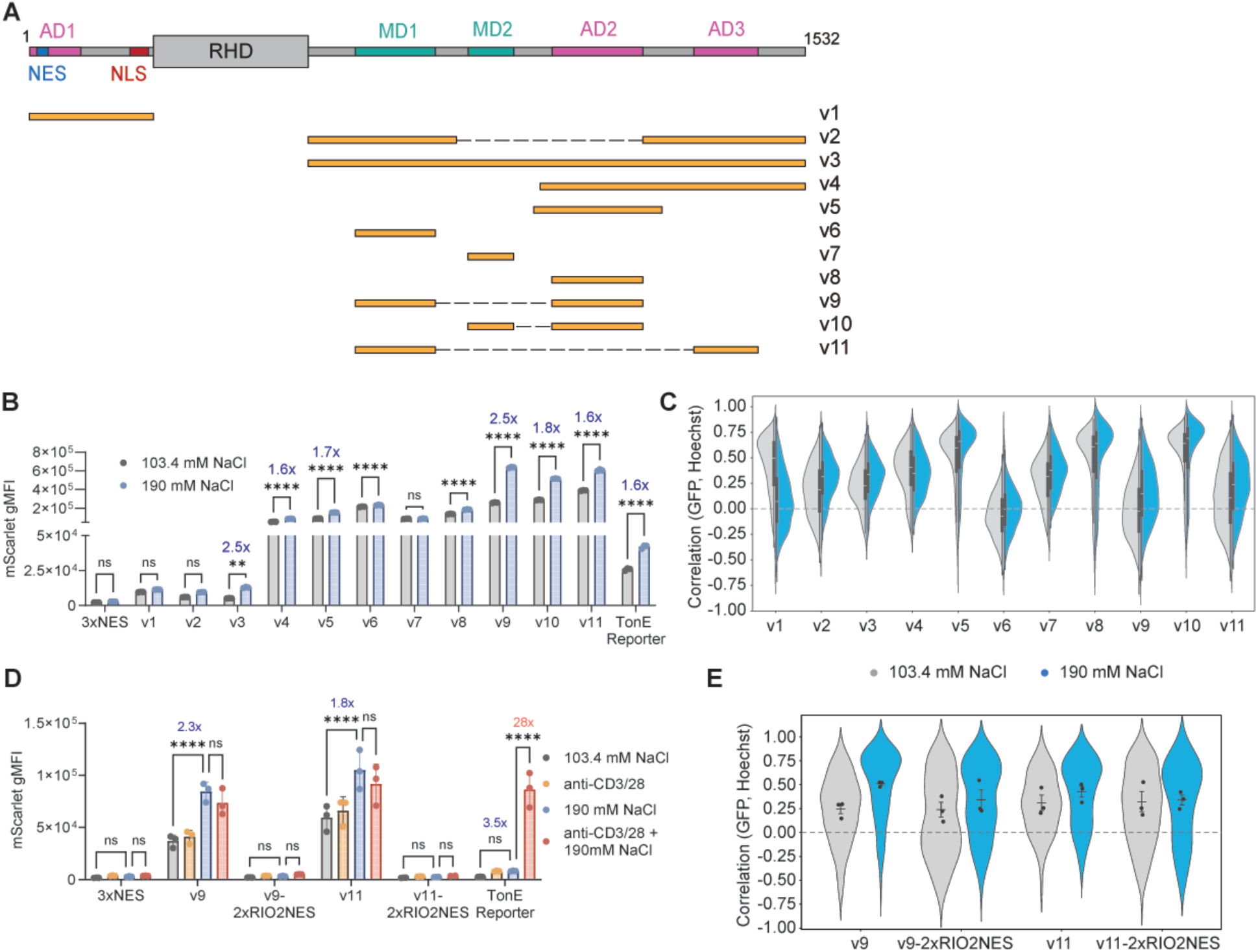
Hyperosmolarity sensing NCtags derived from NFAT5. All panels show cells expressing HEsynTF fused to the indicated NCtags. **A** Domain schematic of NCtags derived from the hyperosmolairty-responsive NFAT5 transcription factor. Orange bars indicatesregions used as the NCtag. **B** Reporter mScarlet gMFI output of GFP⁺ Jurkat cells (190 mM NaCl versus 103 mM NaCl native to RPMI). n = 3 treatment replicates, error bars show SD; significance calculated by two-way ANOVA with Bonferroni’s adjustment for multiple comparisons. **C** Violin plots of single-cell GFP-Hoechst correlation quantified from confocal microscopy images of Jurkat cells after the indicated treatments. Violins represent >150 cells per construct and treatment condition from two combined treatment wells; horizontal line indicates median; black bar for 25-75^th^ quartiles, whiskers for 1.5× the interquartile range. **D** Reporter mScarlet gMFI output of GFP+ primary T cells after the indicated treatments. Fold change shown for 190 mM NaCl versus 103 mM NaCl. n = 3 donors, error bars show SD; significance calculated by two-way ANOVA with Tukey’s adjustment for multiple comparisons. **E** Violin plots of single-cell GFP-Hoechst correlation quantified from confocal microscopy images of primary T cells expressing NFAT5-NCtags after the indicated treatments. Violins represent >120 cells per condition; dots indicate per-donor means (n=3 donors); error bars show SD. All cells are treated for 24 hours (for all panels: ** p < 0.01, *** p < 0.001, **** p < 0.0001).

We next transduced primary T cells with v9 and v11 and treated cells with 190 mM NaCl, anti-CD3/CD28 stimulation, or both, for 24 h. These NCtags responded selectively to hyperosmolarity and not to TCR signaling (Fig. 4D). In contrast, a TonE reporter encoding the response elements for NFAT5 required both signals to induce strong mScarlet. Endogenous NFAT5 is known to be transcriptionally activated by other NFAT-family members after TCR stimulation, explaining this dual dependence on calcium signaling and hyperosmolarity.^32^

Addition of two copies of the RIO2 NES, tested as one route to further reduce background activity, abolished induction upon treatment, indicating that NES-based background reduction is not compatible with all NCtags (Fig. 4D). Nuclear localization of v9 increased more than that of v11 with high salt. However, nuclear translocation for the 2xRIO2-NES versions was modest (Fig. 4E), suggesting that other background-reduction strategies, such as placing the NFAT5-synTF under inducible transcriptional control, as occurs for native NFAT5, offer a more promising route to improve fold change.

### Development of cAMP-responsive NCtags

To demonstrate the modularity of NCtag control over HEsynTF, we next developed NCtags responsive to cAMP signaling. Intracellular cAMP levels are elevated and promote dysfunction in cytotoxic T cells in response to immunosuppressive GPCR ligands present in the tumor microenvironment, including prostaglandin E₂ (PGE₂), adrenaline, and adenosine.^33–36^ We generated NCtags derived from the regulatory regions of human proteins reported to translocate from the cytoplasm to the nucleus in response to cAMP signaling: HDAC4, CRTC2, CREM, and the CREM isoform ICER.^37–39^

Human histone deacetylase 4 (HDAC4) resides predominantly in the cytoplasm but translocates to the nucleus upon activation of the cAMP–PKA signaling pathway.^40,41^ We generated six NCtags incorporating different regulatory regions of HDAC4 (Fig. 5A). Treating transduced Jurkat cells with the cell-permeable cAMP analog dibutyryl-cAMP (dbcAMP) for 24 h induced varying degrees of nuclear translocation across constructs (Fig. 5B). The HDAC4 variants yielded a range of basal reporter activity and cAMP-induced output, with most variants showing modest induction upon dbcAMP treatment (Fig. 5C). v5 achieved the highest dynamic range, with a 7.7-fold increase in reporter output upon dbcAMP treatment relative to DMSO, but showed high basal leakiness. HDAC4v6 emerged as the lead construct, combining low basal activity with ∼5-fold induction upon dbcAMP treatment (Fig. 5B).

**Fig. 5.**
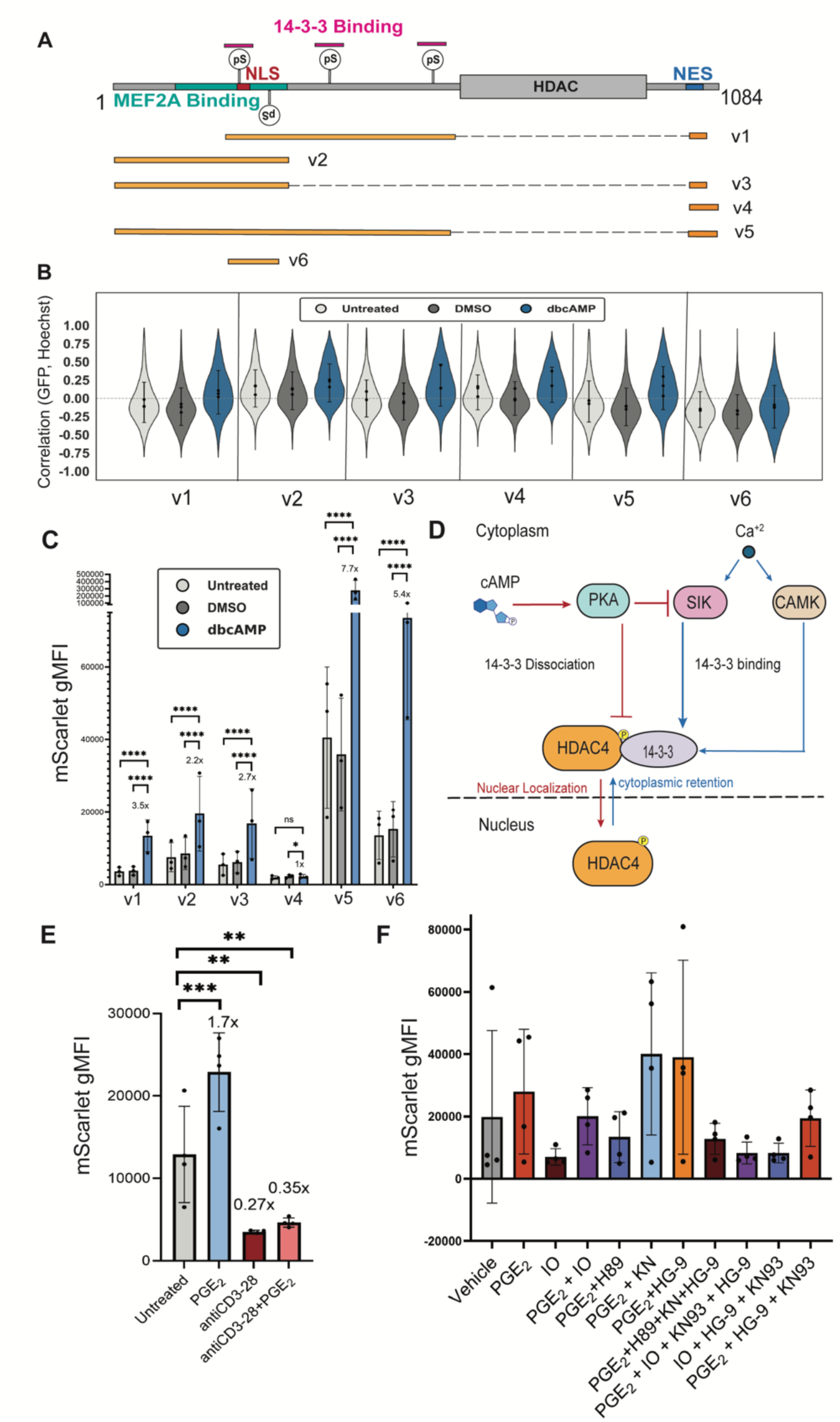
Development of a cAMP-responsive NCtag based on HDAC4. **A** Domain schematic of NCtags derived from the cAMP-responsive HDAC4 protein. Orange bars indicate regions used as the NCtag. Phosphoserines (pS) and regulatory domains are indicated. **B** Violin plots of single-cell GFP-Hoechst correlation quantified from confocal microscopy images of Jurkat cells expressing HDAC4-NCtags fused to HEsynTF after the indicated treatments for 24 hours (500 μM dbcAMP or 1 μM PGE2). Violins represent >2,500 cells per condition; dots indicate per-replicate means (n = 3 treatment replicates); error bars show SD. **C** Reporter mScarlet gMFI output of Jurkat cells expressing HEsynTF fused to HDAC4-NCtags after the indicated treatments for 24 hours (500 μM dbcAMP or 1 μM PGE2). n = 3 treatment replicates, error bars show SD for all panels. Significance across treatment conditions was evaluated using a linear mixed-effects model with experimental day specified as a random intercept, followed by pairwise Wald contrasts with Holm-Bonferroni step-down correction (*p < 0.05,**p < 0.01, ***p < 0.001, ****p < 0.0001, ns = not significant). **D** Schematic of the regulation of HDAC4 subcellular localization by cAMP and Ca²⁺ signaling. cAMP-activated PKA promotes HDAC4 nuclear localization by directly phosphorylating HDAC4 at Ser265/266 and by indirectly inhibiting SIK, both leading to 14-3-3 dissociation and nuclear import. Conversely, Ca²⁺-activated SIK and CaMK phosphorylate HDAC4, promoting 14-3-3 binding and cytoplasmic retention. **E-F** Reporter mScarlet gMFI output of primary T cells expressing HEsynTF fused to the HDAC4v6 NCtag after the indicated treatments for 24 hours (PGE2 (1 μM), anti-CD3/28, ionomycin (IO; 1.34 μM), PKA inhibitor H89 (10 μM), CaMK inhibitor KN-93 (“KN” 5 μM), SIK inhibitor HG-09-91-01 (“HG-9”; 5 μM), vehicle is DMSO, applied alone or in combination as indicated). n = 4 donors; dots indicate per-donor means; error bars show SD. Significance across treatment conditions was evaluated using a linear mixed-effects model with donor identity specified as a random intercept, followed by pairwise Wald contrasts with Holm-Bonferroni step-down correction (*p < 0.05,**p < 0.01, ***p < 0.001, ****p < 0.0001, ns = not significant). For F, all comparisons were ns.

We tried three strategies to further attenuate background activity in v6. The v7 variant, incorporating an endogenous HDAC4 NES (Supplementary Fig. 4A), showed reduced basal activity but also diminished induction (Supplementary Fig. 4B,C). Second, we attempted to reduce synTF expression levels by removing the Woodchuck Hepatitis Virus Posttranscriptional Regulatory Element (WPRE) RNA stabilization element from the construct. This did not reduce the background levels but instead compromised the dynamic range of the tag (Supplementary Fig. 4B,C). Finally, we engineered a variant v8 to include an additional 14-3-3 binding site, because 14-3-3 proteins occur as dimers and two binding sites might increase avidity (Supplementary Fig. 4G). v8 strongly reduced background while maintaining a small portion of cells that translocated in response to dbCAMP, which was accompanied by a very modest increase in reporter induction (Supplementary Fig. 4D-F). HDAC4v6-NCtag therefore remained the optimal candidate for transgene regulation, exhibiting 15-fold induction from dbcAMP in primary T cells or 10-fold induction from the natural cAMP-inducing ligand PGE₂ (Supplementary Fig. 4C).

Phosphorylation of HDAC4 by calcium/calmodulin-dependent protein kinases (CaMKs) or salt-inducible kinases (SIKs) promotes binding to 14-3-3 proteins and cytoplasmic retention, whereas PKA signaling activated by cAMP inhibits SIKs or directly alters HDAC4 phosphorylation to favor nuclear localization (Fig. 5D).^37,40^ To dissect the effect of calcium signaling on HDAC4v6-driven reporter output, we treated primary T cells with PGE₂ or anti-CD3/28, alone or in combination (Fig. 5E). Anti-CD3/28 decreased reporter output compared to the untreated condition and impaired the PGE₂-driven increase in reporter output. To test the contributions of all kinases controlling HDAC4v6, we treated cells with PGE₂ to induce PKA activity, ionomycin to induce CaMK/SIK activity, and the small-molecule inhibitors H89, KN-93, and HG-9-91-01 to inhibit PKA, CaMK, and SIK, respectively (Fig. 5F). Ionomycin decreased baseline and PGE₂-driven reporter output, as observed with anti-CD3/28. KN-93 or HG-9-91-01 treatments increased PGE₂-driven reporter output, while H89 decreased it. These findings demonstrate that the NCtag is regulated by all three kinases, with PKA promoting nuclear translocation and reporter increase, and CaMK and SIK promoting retention in the cytoplasm and decreased reporter output.

To explore NCtags with distinct cAMP and calcium signal integration logic, we next evaluated the CREB-regulated transcription coactivator 2 (CRTC2) protein. We incorporated the regulatory region containing key phosphorylation sites and the NLS and NES sequences as an NCtag (CRTC2v1) fused to HEsynTF (Fig. 6A). Upon transduction into Jurkat cells and 24 hr treatment, we observed an 8-fold increase in mScarlet reporter output with dbcAMP compared to the DMSO control and a 3-fold increase with PGE₂ (Fig. 6B). Notably, ionomycin treatment did not affect reporter output. Localization analysis revealed an increase in nuclear translocation with dbcAMP, whereas PGE₂ and ionomycin correlation values remained similar to baseline levels (Fig. 6C). In primary T cells we observed a similar 2-fold induction of reporter output with both dbcAMP and PGE₂ treatments. Combining PGE₂ and ionomycin resulted in an induction comparable to PGE₂ alone, indicating that calcium signaling does not block nuclear translocation for CRTC2v1 (Fig. 6D). Localization in primary T cells showed increased nuclear translocation with dbcAMP, consistent with Jurkat data, while the PGE₂ + ionomycin condition showed a modestly increased correlation value compared to PGE₂ treatment alone, indicating modest calcium influence on localization that was not reflected in reporter output (Fig. 6E).

**Fig. 6.**
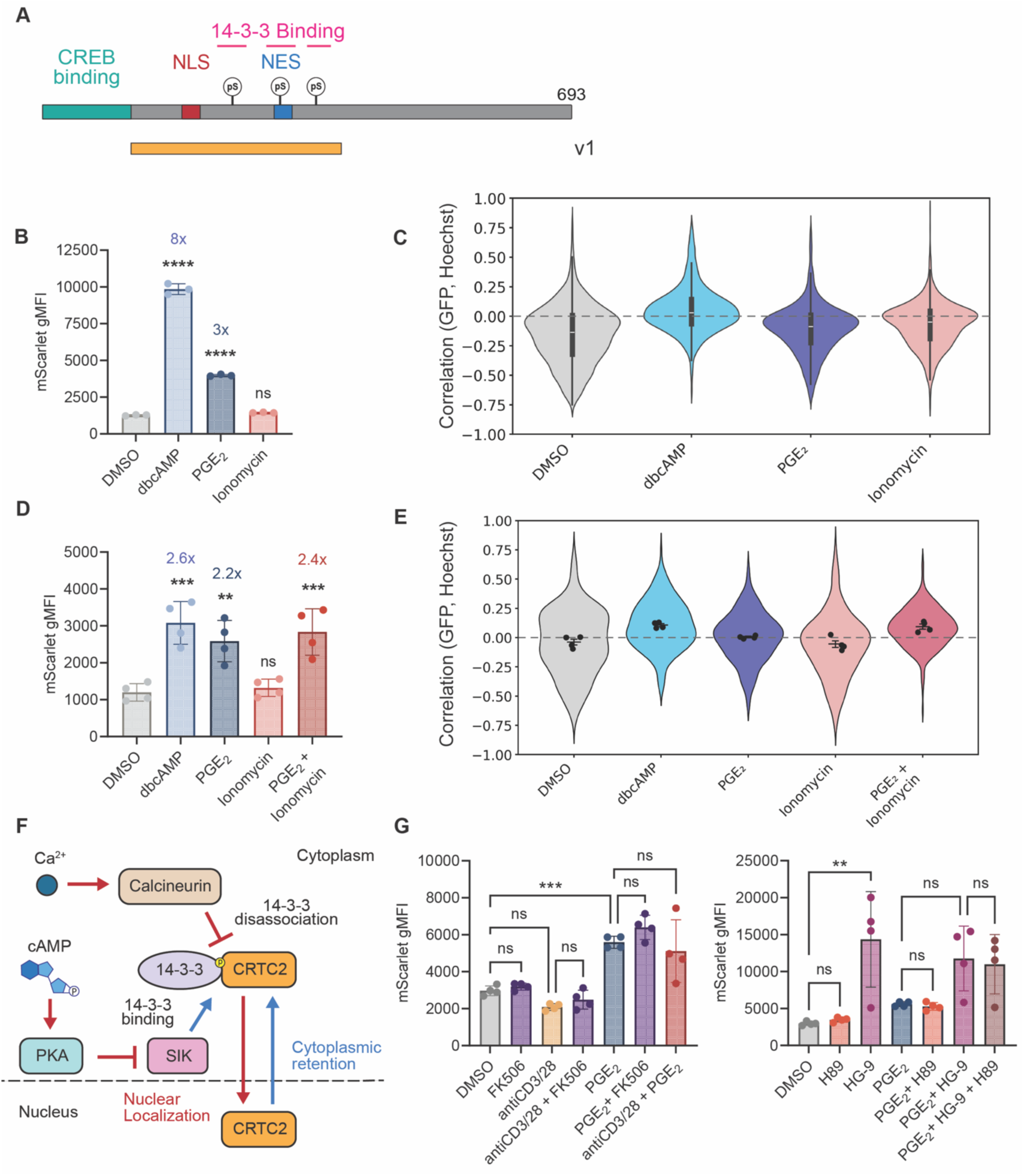
cAMP-responsive NCtag derived from CRTC2. All panels show cells expressing HEsynTF fused to CRTC2v1 NCtag. **A** Domain schematic of NCtag derived from the cAMP-responsive CRTC2 protein. Orange bar indicate region used as the NCtag. **B** Reporter mScarlet gMFI output of GFP⁺ Jurkat cells. n = 3 treatment replicates; error bars show SD; significance calculated by one-way ANOVA with Dunnett’s adjustment for multiple comparison vs. DMSO. **C** Single-cell correlation of GFP and Hoechst from confocal microscopy of Jurkat cells. Violins represent >1,500 cells per condition from three combined treatment wells; horizontal line indicates median; black bar for 25-75^th^ quartiles; whiskers for 1.5× the interquartile range. **D** Reporter mScarlet gMFI output of GFP⁺ primary T cells. n = 4 donors; error bars show SD; significance calculated by one-way ANOVA with Dunnett’s adjustment for multiple comparison vs. DMSO. **E** Single-cell correlation of GFP and Hoechst from confocal microscopy of primary T cells. Violins represent >250 cells per condition; dots indicate per-donor means (n = 4 donors); error bars show SD. **F** Schematic of the regulation of CRTC2 subcellular localization. Under basal conditions, CRTC2 is phosphorylated by SIK and retained in the cytoplasm via interactions with 14-3-3. Both cAMP and calcium signals promote CRTC2 dephosphorylation, through SIK inactivation by PKA and calcineurin activation, respectively, releasing CRTC2 from 14-3-3 and enabling nuclear translocation. Red arrows indicate nuclear localization; blue arrows indicate cytoplasmic retention. Red arrows indicate nuclear localization promotion and red arrow indicate cytoplasmic rentention promotion. **G-H** Reporter mScarlet gMFI output of GFP⁺ primary T cells. Treatments applied alone or in combination as indicated: PGE_2_ (1 µM), anti-CD3/28, ionomycin (1.34 µM), H89 (10 µM), FK506 (5 µM), HG-09-91-01 (“HG-9”; 5 µM). n = 4 donors; dots indicate per-donor means; error bars show SD; significance calculated by one-way ANOVA with Tukey’s adjustment for multiple comparisons. All cells are treated for 24 hours (for all panels: ** p < 0.01, *** p < 0.001, **** p < 0.0001).

CRTC2 is retained in the cytoplasm through phosphorylation by SIKs promoting interactions with 14-3-3 proteins. cAMP and calcium signals promote CRTC2 dephosphorylation through inhibition of SIKs by PKA and activation of calcineurin, respectively (Fig. 6F).^38^ To further dissect the contribution of calcium to CRTC2v1-driven reporter output, we treated primary T cells with PGE₂, anti-CD3/28, or the calcineurin inhibitor FK506, alone or in combination (Fig. 6G). PGE₂ increased reporter output with or without anti-CD3/28, while anti-CD3/28 alone or in combination with FK506 had no effect on reporter output. These results indicate that calcineurin activation by ionomycin or anti-CD3/28 does not affect PGE₂-driven increase of CRTC2 nuclear localization or reporter output. To examine the involvement of PKA and SIK in cAMP-driven reporter output, we treated with H89 or HG-9-91-01 alone or in combination with PGE₂ (Fig. 6H). HG-9-91-01 increased reporter output, indicating that SIK activity retains CRTC2 in the cytoplasm. However, H89 did not decrease reporter output, suggesting that kinases other than PKA might contribute to SIK inhibition and CRTC2 nuclear localization and reporter output.

To test additional putative cAMP-responsive NCtags, we generated two constructs from CREM and one from ICERII, an isoform of CREM, which are reported to translocate in response to cAMP signaling (Supplementary Fig. 5A,B).^39,42^ The NCtags were transduced in Jurkat cells and dbCAMP treatment led to substantial nuclear translocation for the tag derived from CREM KID domain (CREM-KID) and no change in localization for the CREM NCtag with the deleted leucine zipper (CREMdLZ) (Supplementary Fig. 5C,D). The NCtag derived from ICERII without the leucine zipper (ICERIIdLZ) also showed moderate nuclear translocation (Supplementary Fig. 5B,E). All three tags showed a 2-fold increase in reporter output in response to dbCAMP (Supplementary Fig. 5F-H). Together, these results show that regulatory regions from multiple localization-regulated proteins can serve as cAMP-inducible NCtags capable of HEsynTF control.

### NCtags respond to endogenous cAMP and calcium signals with distinct logic behaviors in tumor spheroids

We next sought to directly compare how cAMP and calcium signals are integrated by the NFAT1rr, HDAC4v6, and CRTC2v1 NCtags. We first treated primary T cells with dbcAMP, anti-CD3/28, or both (Fig. 7A,B). NFAT1rr showed increased reporter output with anti-CD3/28 alone; however, addition of dbcAMP modestly reduced this response (a partial NIMPLY gate with cAMP inhibiting calcium-driven output), consistent with evidence that cAMP/PKA signaling phosphorylates NFATs to promote 14-3-3 binding and cytoplasmic retention.^43^ HDAC4v6 showed increased reporter output when both treatments were combined or with dbcAMP alone (acting as an IMPLY gate), in contrast to what was observed previously for combination of ionomycin or anti-CD3/CD28 with PGE₂, where it behaved more like a NIMPLY gate with calcium inhibiting cAMP-driven output (Fig. 5F,G). For CRTC2v1, stimulation with anti-CD3/28 and dbcAMP potentiated the reporter output increase from dbcAMP alone, a response also not observed with PGE₂, where anti-CD3/CD28 had no effect (Fig. 6G). Thus, on CRTC2v1, calcium and cAMP signaling form an IMPLY gate with calcium having no effect on cAMP-induced output, or calcium can further increase output, depending on the stimuli used.

**Fig. 7.**
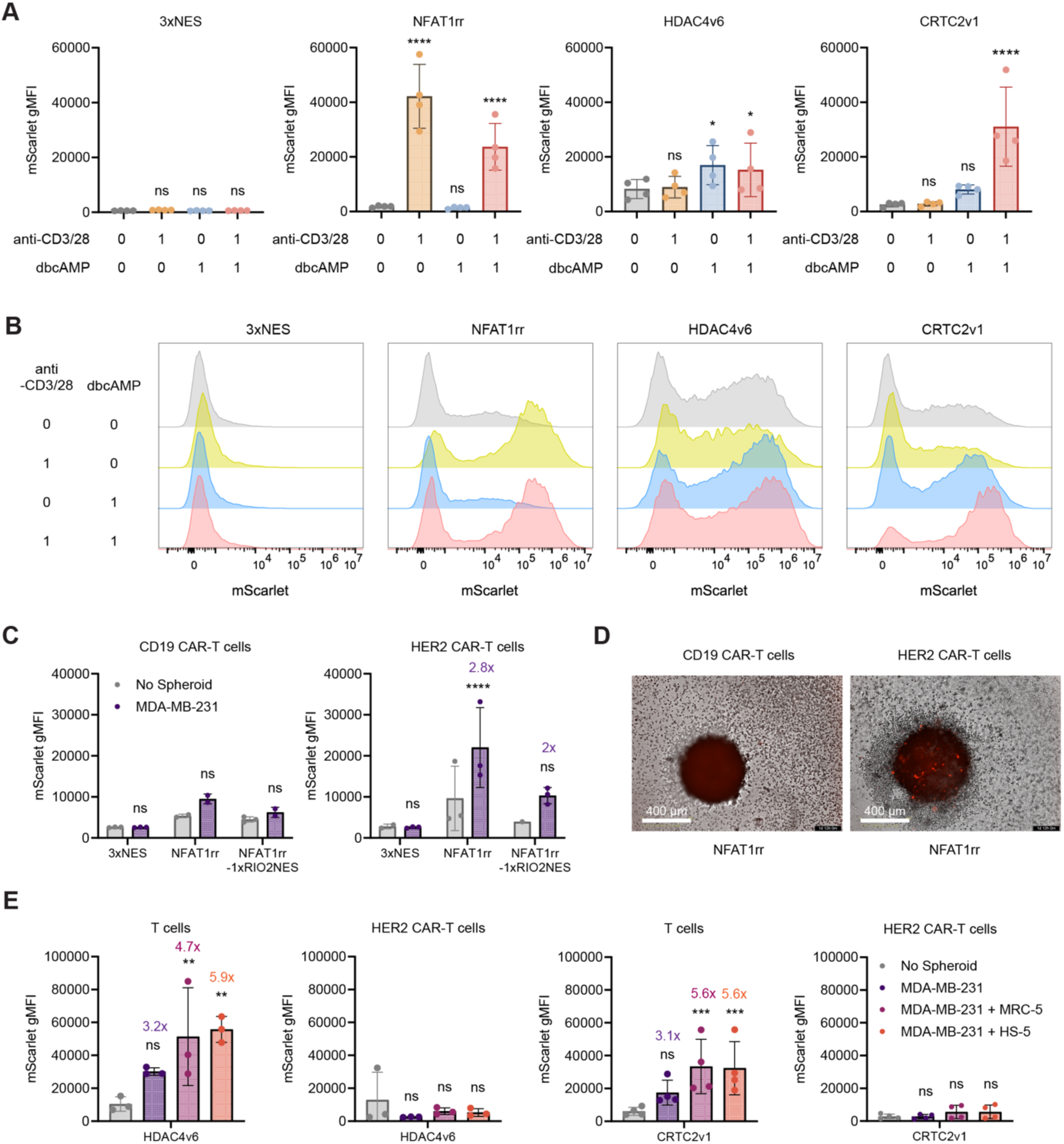
NCtags integrate cAMP and calcium signals with distinct logic behaviors. All panels show primary T cells expressing HEsynTF fused to the indicated NCtags, unless otherwise noted. **A** Reporter mScarlet gMFI output of GFP⁺ cells after treatment with plate-bound anti-CD3/28, dbcAMP, or a combination of both for 24 hours. n = 4 donors; error bars show SD; significance calculated by linear mixed-effects model with NCtag × Treatment fixed effects and donor as a random intercept; pairwise comparisons vs. no treatment evaluated by Wald test. **B** Representative flow cytometry histograms depicting mScarlet reporter output of GFP⁺ primary T cells of one donor under the indicated treatments. **C** Reporter mScarlet gMFI output in CD19⁺ or HER2 CAR ⁺ GFP⁺ T cells after 72 hours of co-culture with MDA-MB-231 tumor spheroids. n = 3 donors; error bars show SD; significance calculated by mixed-effects analysis with Bonferroni’s adjustment for multiple comparisons vs. no spheroid condition. **D** Representative images of MDA-MB-231 spheroids after 36 hours of co-culture with CD19 or HER2 CAR-T cells expressing depicting mScarlet reporter output of one donor. **E** Reporter mScarlet gMFI output of GFP ⁺ primary T cells or HER2 CAR⁺ GFP⁺ T cell safter 62 hours of co-culture with MDA-MB-231 and HS-5 or MRC-5 tumor spheroids. n = 3 biological replicates (HDAC4v6), n = 4 biological replicates (CRTC2v1); error bars show SD; significance calculated by two-way ANOVA with Dunnett’s adjustment for comparisons vs. no spheroid condition. All cells are treated for 24 hours, unless otherwise noted (for all panels: * p < 0.05, ** p < 0.01, *** p < 0.001, **** p < 0.0001).

We assessed the temporal reversibility of NCtags by treating cells with anti-CD3/28 (NFAT1rr) or dbcAMP (HDAC4v6 and CRTC2v1) for 24 hours, removing the stimulus and replacing with normal RPMI for 72 hours, and comparing reporter output to continuously treated or untreated cells for 96 hours. This demonstrated the ability of NCtags to turn off transgene expression upon signal withdrawal (Supplementary Fig. 6A, B).

We next examined whether NFAT1rr, HDAC4v6, and CRTC2v1 NCtags could respond to endogenous cAMP and calcium signals from HER2⁺ MDA-MB-231 tumor spheroids. We first quantified tumor-derived PGE₂ by ELISA and found that tumor spheroids alone produced approximately 1,000 pg/mL PGE₂, which increased to 1,500 pg/mL upon addition of cancer-associated fibroblasts, which are known to produce high levels of PGE₂ (Supplementary Fig. 7A).^44,45^ To provide antigen-driven calcium signaling, primary T cells were transduced with a HER2 CAR following transduction with the three HEsynTF-NCtag constructs; for NFAT1rr, a CD19 CAR was also used as a non-cognate control, while HDAC4v6 and CRTC2v1 cells were tested without a CAR to isolate responses to PGE₂. T cells were cultured with tumor spheroids for 72 hours and reporter output was analyzed by flow cytometry on disrupted spheroids. For NFAT1rr, only HER2 CAR-T cells and not CD19 CAR-T cells showed increased reporter output, demonstrating that HEsynTF-NFAT1rr can respond specifically to antigen-driven signaling in a tumor co-culture context (Fig. 7C, D). In contrast, CRTC2v1 and HDAC4v6 T cells without CAR showed increased reporter output after co-culture with tumor spheroids formed from MDA-MB-231 alone or with HS-5 or MRC-5 fibroblasts, indicating responsiveness to tumor-derived PGE₂ (Fig. 7E). However, HER2 CAR-T cells expressing HDAC4v6 or CRTC2v1 did not show increased reporter output, consistent with our earlier observation that calcium signaling suppresses HDAC4v6 when combined with PGE₂ (Fig. 5F, G) (NIMPLY gate), and indicating that CAR-driven signaling is sufficient to counteract PGE₂-driven CRTC2v1 induction in this context (NIMPLY gate), unlike what we observed with ionomycin or anti-CD3/28 treatments (Fig. 6D, 6G, 7A).

Finally, as a proof of concept for the application of the HEsynTF-NCtag platform to control therapeutic transgenes, we used HDAC4v6 to control HER2 CAR expression in response to physiological levels of PGE₂. A PGE₂ titration in T cells expressing HEsynTF-HDAC4v6 confirmed mScarlet reporter output at concentrations measured in MDA-MB-231 tumor spheroid supernatant (Supplementary Fig. 7B). We next replaced mScarlet with the HER2 CAR transgene and observed increased CAR expression upon dbcAMP treatment, while T cells expressing HER2 CAR under the HEsynTF-3xNES control or the constitutive EFS promoter did not significantly change CAR expression (Supplementary Fig. 7C). After co-culture with MDA-MB-231-GFP/MRC-5 spheroids, we observed a significant 1.8-fold increase in HER2 CAR expression in T cells expressing HDAC4v6 compared to controls (Supplementary Fig. 7D). We also observed that NCtag-driven HER2 CAR induction produced tumor killing comparable to constitutively expressed HER2 CAR (Supplementary Fig. 7E, F), demonstrating a functional output of and NCtag-synTF driven by endogenous stimuli.

## Discussion

We developed a modular platform for sense-and-response to intracellular signaling based on post-translational control of synthetic TF localization. By pairing NCtags with a synTF, we generated genetic circuits responsive to calcium signaling, hyperosmolarity, and cAMP/PKA signaling in T cells. We also identified properties of synTFs that enable effective regulation at the level of nucleocytoplasmic localization, optimized an all-in-one lentiviral architecture for delivery into primary human T cells, and demonstrated that NCtags can be isolated from the regulatory regions of multiple human proteins.

Intracellular sensing with NCtags provides a complementary approach to synthetic receptor systems. Receptor-based circuits typically detect a defined extracellular ligand or cell-surface antigen,^8,12^ whereas intracellular signaling pathways can integrate information from multiple receptor-initiated or metabolic signaling pathways. We demonstrated this concept using NFAT1rr, HDAC4v6 and CRTC2v1 NCtags in breast cancer spheroids, which produced reporter output depending on logical combinations of calcium (from CAR-antigen) and cAMP (from tumor-supplied PGE_2_) signaling. While each NCtag displayed roughly IMPLY or NIMPLY logic behaviors depending on the source of combination of cAMP and calcium inputs, the specific output level is expected to vary across cell types and contexts depending on the balance of each second messenger. In the tumor spheroid context, for example, molecules other than PGE₂ may also contribute to cAMP signaling, further modulating the output.

NCtags enable engineered cells to respond to the downstream effects of combinations of environmental inputs, allowing them to sense a cellular signaling state rather than a single extracellular molecule and providing a general framework for cells to dynamically adapt their behavior to their environments. The inputs we chose are themselves therapeutically motivated: PGE_2_, adenosine, and other tumor microenvironment factors elevate cAMP in cytotoxic T cells, macrophages, and NK cells and suppress their function,^34,36,46,47^ and hyperosmolarity is a feature of necrotic tumor cores and inflamed tissues.^48,49^ NCtags built on these pathways will enable conversion of signals that normally drive dysfunction into an inducible therapeutic output.

The NCtag-synTF platform has advantages in modularity relative to promoters controlled by signaling-inducible native TFs. Native TF-responsive promoters couple transgene expression to the abundance, DNA-binding activity, and transcriptional activation strength of endogenous TFs. These properties are difficult to alter in a modular fashion and place constraints on the maximum expression level of the transgene, as has been noted in the induction of CAR transgenes under the control of ATF4 or HIF-1*a*.^50,51^ In contrast, isolating signal sensing allows NCtags to be combined with a synTF whose transcriptional properties can be engineered for the desired output. While most of our applications used the HEsynTF composed of HNF1A DBD and en3x9 TAD, we demonstrated that the stronger p65 TAD may also be suitable for NCtag control, with appropriate synTF expression tuning to reduce background. Likewise, we showed that other homeodomains like PAX6 are amenable to localization control, providing options for cell or tissue applications where HNF1A may be insufficiently orthogonal. These choices reflect our translation-focused design. The platform is built from human-derived DBDs, TADs, and native regulatory regions, and its compact footprint fits alongside the transgene in a single clinically relevant vector, providing advantages in predicted immunogenicity and manufacturing relative to TALE- or dCas9-based systems.^1,13,17^

The modularity of the NCtag-synTF platform may also facilitate future inclusion in more complex synthetic biology circuits in which NCtag-synTF expression is inducible by another stimulus. Native TFs often layer transcriptional and post-translational regulation, as demonstrated by the dual requirement for calcium-dependent transcriptional induction plus post-translational hyperosmolarity induction of the TonE reporter by NFAT5 (Fig. 4D).^31,52^ Layering NCtag-synTF control with inducible expression of the NCtag-synTF would allow combined transcriptional and post-translational AND-gates, and may reduce background from some NCtags like NFAT5v9/v11. We demonstrated the ability to tune background induction levels by adding additional cytoplasmic localization sequences (NES or 14-3-3 binding sites) to NFAT1rr and HDAC4v6 (Fig. 3H, Supplementary Fig. 4), providing options for transgenes applications with different levels of tolerance for background expression.

Although we apply NCtags here to control a synTF, they could potentially be adapted to regulate other classes of proteins whose functions depend on subcellular localization, including native TFs or epigenetic regulators, or signaling proteins only active in the cytoplasm. NCtag-synTFs may also serve as integrating biosensors. The cAMP/PKA biosensor field has largely produced fast fluorescent or bioluminescent reporters optimized for microscopy, which report signaling but cannot actuate an output.^18,53^ An NCtag-synTF instead converts a signaling state into a durable, genetically encoded readout or functional transgene.

NCtag localization is governed by the relative binding affinities to importins, exportins, and cytosolic anchor proteins like 14-3-3. The sequence-to-localization rules for NCtags must be better defined before we can predict or design such regulatory regions. Design heuristics have been developed for KTRs,^18,54^ which generally translocate from the nucleus to cytoplasm in response to phosphorylation, but equivalent rules for cytoplasm-to-nucleus translocating NCtags remain undefined. Our HDAC4 and CRTC2 NCtags exploit a natural architecture of overlapping 14-3-3 sites and NLS that integrates SIK and PKA activity. Designing such regions *de novo* would require signaling knowledge beyond sequence-to-binding relationships. A related limitation is that our NCtags are derived from natural regulatory regions, so we cannot exclude their regulation by additional signaling inputs. Minimization of these regions, or development of fully synthetic phospho-regulated NLS/NES elements, should yield more pathway-specific NCtags and more predictable behavior.

Together, this work establishes nucleocytoplasmic localization control as a general and modular route to sense-and-response inside engineered cells. By separating signal sensing, encoded in an exchangeable NCtag, from transcriptional output, encoded in an engineerable human synTF, the platform links intracellular signaling states to programmable transgene expression. Implemented with human-derived components in primary human T cells and delivered in a single lentiviral vector, the platform is positioned for *in vivo* evaluation to determine the spatiotemporal specificity of NCtag-driven transgene induction in a native tumor microenvironment. We anticipate that NCtags will provide an extensible foundation for therapeutic cells that read their own signaling state and respond accordingly.

## Methods

### Vector design and molecular cloning

Synthetic gene fragments encoding all components were synthesized by Twist Bioscience and cloned into either pLenti (lentiviral) or SFG (gamma-retroviral) backbones using Gibson Assembly with NEBuilder HiFi Assembly Master Mix (New England Biolabs). Constructs were transformed into chemically competent *Escherichia coli* DH5α cells and selected on antibiotic-containing LB agar plates. Plasmid DNA was purified using Zymo Miniprep or Qiagen Maxiprep kits. All vectors were sequence-verified by Sanger sequencing or full-plasmid nanopore sequencing prior to use.

All protein sequences are provided in Supplementary Table 1. Human TF sequences were sourced from the canonical UniProt entry. For CREM, which has many isoforms, we used CREM-BCEFGgammaHIbeta, Uniprot Q03060-5, and unless otherwise specified, nuclear localization signals (NLS) and nuclear export signals (NES) correspond to a 2xSV40-derived NLS plus a nucleoplasmin NLS and three copies of an NES derived from HIV-Rev, respectively. The membrane-targeting CAAX motif was derived from human KRAS4B. SynTF architectures consisted of, from N-to-C, DBD-TAD-mGreenLantern-NCtag. Reporter-only constructs were cloned to express a constitutive tNGFR surface marker for detection of transduced cells. The number and identity of response elements used for synTF-responsive reporters in Figure 2 are detailed in Supplementary Table 2, and an example all-in-one lentiviral sequence is provided in Supplementary Table 3. Plasmids used for each figure panel are listed in Supplementary Table 4, and full sequences are available upon request. The synthetic minimal promoter used has been reported in prior inducible cell therapy constructs.^20^ The HER2.28z CAR sequence used the FRP5 scFV, IgG1 short hinge, and CD28 costimulatory domain.^55^ The CD19.28z CAR sequence used the FMC63 scFV, a FLAG epitope marker, IgG2 short hinge, and CD28 costimulatory domain.^56^

### Cell isolation and culture

Peripheral blood mononuclear cells (PBMCs) were isolated from healthy donor buffy coats or leukocyte reduction chambers obtained from the Gulf Coast Regional Blood Center (Houston, TX, USA) by density gradient centrifugation using Lymphoprep™ (STEMCELL Technologies). All cells were maintained at 37 °C in a humidified incubator with 5% CO₂.

Primary human T cells and Jurkat cells (gift from Maksim Mamonkin, verified by short tandem repeat (STR) sequencing) were cultured in RPMI-1640 supplemented with 10% heat-inactivated fetal bovine serum (FBS, Gibco) and 2 mM GlutaMAX (Gibco). For expansion of primary T cells, recombinant human IL-7 and IL-15 were added at 10 ng/mL every 2 or 3 days.

HEK293T cells used for viral production were cultured in DMEM or IMDM supplemented with 10% FBS and 2 mM GlutaMAX. Breast cancer cells (MDA-MB-231) (gift from Valentina Hoyos, verified by STR sequencing) and stromal cells (Hs5, MRC-5, and primary human breast cancer-associated fibroblasts, gifts from Mary Katherine McKenna, immortalized lines verified by STR sequencing) were maintained in DMEM supplemented with 10% FBS and 2 mM GlutaMAX.

### Viral production and transduction

Retroviral vectors were produced by transient transfection of HEK293T cells. Cells were co-transfected with the transfer plasmid and packaging plasmids according to established protocols.^57^ Transfections were performed using GeneJuice (Novagen) following the manufacturer’s instructions. Viral supernatants were harvested 48-72 h post-transfection and filtered through 0.45 µm membranes.

To achieve higher titer virus for primary T cell transduction, lentiviral vectors were produced by transfecting 4.5x10^6^ CRISPRed HEK293T to Disrupt Antiviral Response cells (CHEDAR) (gift from Donald Kohn)^58^ cells in a 10 cm dish with the AIO lentiviral transfer vector, the second-generation packaging plasmid psPAX2 (Addgene #12260), and the RD114A envelope plasmid^59^ in 10 mL of DMEM. After 6 h, media was replaced with fresh DMEM supplemented with 20 µL of ViralBoost (Alstem). Viral supernatants were collected at 48 h post-replacement and filtered through a 0.45 µm membrane. 24-well non-tissue-culture-treated plates were coated with RetroNectin (Takara) at 7 µg/mL overnight, after which 2 mL of viral supernatant was added to each well and centrifuged for 2 h at 2,000 × g at 32°C.

T cells were transduced at a density of 500,000 cells per well in 2 mL of RPMI according to established protocols.^57^ Jurkat cells were thawed and cultured for 48 h prior to transduction. PBMCs were thawed and stimulated for 48 h with plate-bound anti-CD3 (OKT3, 1 µg/mL) and anti-CD28 (1 µg/mL) antibodies prior to transduction. For dual-modified cells, a sequential transduction strategy was employed. In Fig. 2 and Fig. 3B–F, cells were first transduced with the retroviral synTF-driven mScarlet reporter vector, followed by the indicated retroviral synTF(-NCtag) construct vectors 24 h later. In Fig. 7C, D, cells were first transduced with the indicated AIO lentiviral HEsynTF-NCtag vector, followed by the retroviral CAR vector 24 h later.

### Treatments and stimulation

T-cell receptor stimulation was performed using plate-bound anti-CD3 (OKT3) and anti-CD28 antibodies at a final coating concentration of 1 µg/mL each in 0.5 mL of water per well of a 24-well non-tissue-culture-treated plate. Plates were incubated overnight at 4°C prior to use. eBioscience™ Cell Stimulation Cocktail (PMA/IO; Invitrogen) was used at 1× final concentration, ionomycin (StemCell Technologies) was used at a final concentration of 1.34 µM, dibutyryl-cAMP (dbcAMP; Cayman Chemical) was used at a final concentration of 500 µM and prostaglandin E₂ (PGE₂; Cayman Chemical) at a final concentration of 1 µM, unless otherwise noted. Osmotic control treatments were performed using a final concentration of 190 mM NaCl in RPMI, or the baseline media level of 103 mM NaCl.

### Flow cytometry

mScarlet reporter expression and mGreenLantern-tagged NCtag construct expression were analyzed using a CytoFLEX LX flow cytometer. HER2 CAR expression was assessed using an Alexa Fluor 647-conjugated AffiniPure Rabbit Anti-Mouse IgG F(ab’)₂ fragment specific antibody (Jackson ImmunoResearch Laboratories). CD19 CAR expression was assessed using a PE/Cyanine7-conjugated anti-DYKDDDDK (FLAG) tag antibody (BioLegend). Expression of reporter-only constructs was assesed using PerCP/Cyanine5.5 anti-human CD271 (NGFR) antibody (BioLegend).

Dead cells were excluded using eFluor 450 viability dye. Data were analyzed using FlowJo v10. For graphing of mScarlet reporter output, gMFI was obtained from GFP⁺ cells expressing a synTF or an NCtag–synTF fusion construct, or from tNGFR⁺ cells expressing a reporter-only construct. For dual-modified cells in Figs. 2B, D and 3E–F, cells were gated first on tNGFR⁺, then on GFP⁺. In Fig. 7C, D, cells were gated first on CD19 CAR⁺ or HER2 CAR⁺, then on GFP⁺. For experiments completed on separate days and graphed together, mScarlet gMFI was normalized to the background gMFI of untransduced cells gated on live cells and analyzed on each day.

### Confocal microscopy

For intracellular localization studies, treated cells were stained with Hoechst 33342 (Thermo Fisher Scientific), fixed with 4% paraformaldehyde for 15 mins onto 96-well glass-bottom plates pre-coated with 12.5 µg/mL fibronectin.^60^ Imaging was performed on a Yokogawa CV8000 spinning disk confocal microscope with a 40x water-immersion lens.

### Image quantification

Automated image analysis was performed using CellProfiler.^61^ Nuclei were segmented based on Hoechst staining, followed by cytoplasmic segmentation using a nuclear mask expansion (8 pixels). Nucleocytoplasmic localization was quantified using the MeasureColocalization module to calculate correlation coefficients between GFP fluorescence and Hoechst signals. A custom Python pipeline was used for downstream filtering to exclude untransduced cells and non-cellular debris based on size and eccentricity thresholds. Representative images were prepared using ImageJ.^62^

### 3D spheroid assay and PGE₂ quantification

50,000 MDA-MB-231 cancer cells were embedded in 2.5% Matrigel and seeded into flat-bottom 96-well plates pre-coated with 1% agarose to prevent adhesion. In experiments incorporating fibroblasts, 12,500 fibroblasts were co-seeded with 37,500 MDA-MB-231 cells (1:3 fibroblast-to-tumor ratio) without Matrigel. For PGE₂ quantification, supernatants were collected after 3 days of spheroid maturation, and PGE₂ was quantified using a competitive ELISA kit (Cayman Chemical) according to the manufacturer’s instructions.

For T cell–spheroid co-culture experiments, T cells were seeded at a 2:1 effector-to-target ratio after 3 days of spheroid formation, normalized to the frequency of GFP⁺ NCtag-synTF-transduced cells; for CAR-T cell experiments, normalization was based on CAR⁺ cell frequency. During co-culture, 96-well plates were imaged every 2 hours using an Incucyte SX5 or CX3 live-cell imaging system (Sartorius) to track mScarlet reporter expression and tumor spheroid GFP integrated intensity over time as a measure of tumor killing. After the indicated co-culture duration, T cells were harvested from spheroids for flow cytometry analysis.

### Statistical analysis

Statistical analyses were performed using GraphPad Prism v10. Data are presented as mean ± SD. ANOVA with an appropriate post-hoc test was used with p < 0.05 considered statistically significant.

Quantification of nucleocytoplasmic localization was evaluated using linear mixed-effects models (LMM) fitted via Restricted Maximum Likelihood (REML) in Python (statsmodels v0.14) to account for hierarchical biological data structure and donor-to-donor variability across independent human donors or biological replicates. Fixed effects were specified for different conditions and constructs. Random intercepts were assigned to individual T-cell donors to control inter-donor baseline variance, with single-cell observations nested within donors. To capture both single-cell response heterogeneity and inter-donor biological reproducibility, statistical testing was conducted at two levels: 1) Single-Cell Population Level: Contrast testing across individual cells evaluated broad population translocation dynamics, reported using standardized z-statistics. 2) Donor-Aggregated Level: Contrast testing across biological replicates evaluated the consistency of construct response across donors, reported using t-statistics with donor-number degrees of freedom. Post-hoc pairwise contrasts assessed significant changes in localization relative to the t = 0 baseline (Figure 1, Figure 3B) or control/untreated conditions (Figs. 4E, 5B, 6E). Fixed-effect size estimates are reported as coefficients (Beta) with standard errors (Std.Err). p-values were adjusted for multiple comparisons using the Holm-Bonferroni sequential step-down method, with statistical significance defined at p < 0.05.

For datasets without biological replicates (Figs. 2B,D, 4C, 6C), an ordinary least squares (OLS) model was used instead, with analysis conducted at the single-cell level only and reported using z-statistics. Fixed-effect size estimates are reported as coefficients (Beta) with standard errors (SE). p-values were adjusted for multiple comparisons using the Holm-Bonferroni correction, with statistical significance defined at p < 0.05.

Flow cytometry analysis was evaluated using LMM implemented in Python (statsmodels v0.14) and GraphPad Prism (v10). For primary human T-cell cohorts (N = 3 donors) and multi-day Jurkat assays (N = 3 independent days), data were modeled using linear mixed-effects models (LMM) fitted via Restricted Maximum Likelihood (REML), specifying treatment conditions and construct variants as fixed effects and donor or experimental day as a random intercept to account for baseline shifts. For single-day Jurkat characterization assays conducted in technical replicate tubes (n = 3), differences across conditions were analyzed using ordinary one-way ANOVA. Pairwise contrasts between vehicle and treatment conditions were evaluated using Wald tests (LMM) or two-tailed Student’s t-tests based on pooled model residuals (ANOVA). Family-wise error rates were controlled within each construct family using the Holm–Bonferroni sequential step-down adjustment, with statistical significance defined at p < 0.05 (*p < 0.05,p < 0.01, ***p < 0.001, ****p < 0.0001, ns = not significant).

Full outputs of statistical tests are given for each figure in Supplementary Table 5.

### Immunogenicity prediction

Predicted MHC class I immunogenicity was computed using NetMHCpan 4.1 EL across 27 HLA-A and HLA-B alleles.^63^ For each overlapping 9-mer peptide, the minimum predicted percentile rank across alleles and the number of alleles with percentile <2 (likely immunogenic epitopes) were determined. To visualize the immunogenic potential of non-human peptides, an immunogenic burden score per peptide was calculated as the product of non-human status (0,1), binding strength (−log10 of the minimum percentile), and allele coverage (fraction of alleles with percentile <2).

## Data availability

Raw data generated in this study are included in the Source Data file. A full list of sequences can be found in the Supplementary Information section. Source data are provided with this paper.

## Acknowledgements

This project was supported by Baylor College of Medicine’s Integrated Microscopy Core with funding from the NIH (P30 DK056338, P30 CA125123, S10OD030414). This project was supported by the Cytometry and Cell Sorting Core at Baylor College of Medicine with funding from the CPRIT Core Facility Support Award (CPRIT-RP240432) and the NIH (CA125123, OD036336, and OD038251), and the assistance of Joel M. Sederstrom. We acknowledge a Texas Childrens’ Hospital Research Institute instrument grant for a Celleca cell counter used in this work. IncuCyte imaging was supported by NIH S10 Grant 1S10OD036406 and performed in the Center for Cell and Gene Therapy at Baylor College of Medicine.

## Author contributions

P.G.D., B.B., and G.F. conceived of the work and designed the experiments. P.G.D., B.B., M.T.M., K.K., and A.O. designed and cloned the constructs. P.G.D., B.B., M.T.M., K.K., and A.O. conducted experiments and analyzed data. P.G.D. and B.B. developed the image quantification and analysis workflow and wrote the associated code. P.G.D. and B.B. made figures. P.G.D., B.B., and G.F. wrote the manuscript. G.F. supervised the work and acquired funding. All authors reviewed and approved the final manuscript.

## Supplementary Figures

**Supplementary Figure 1.**
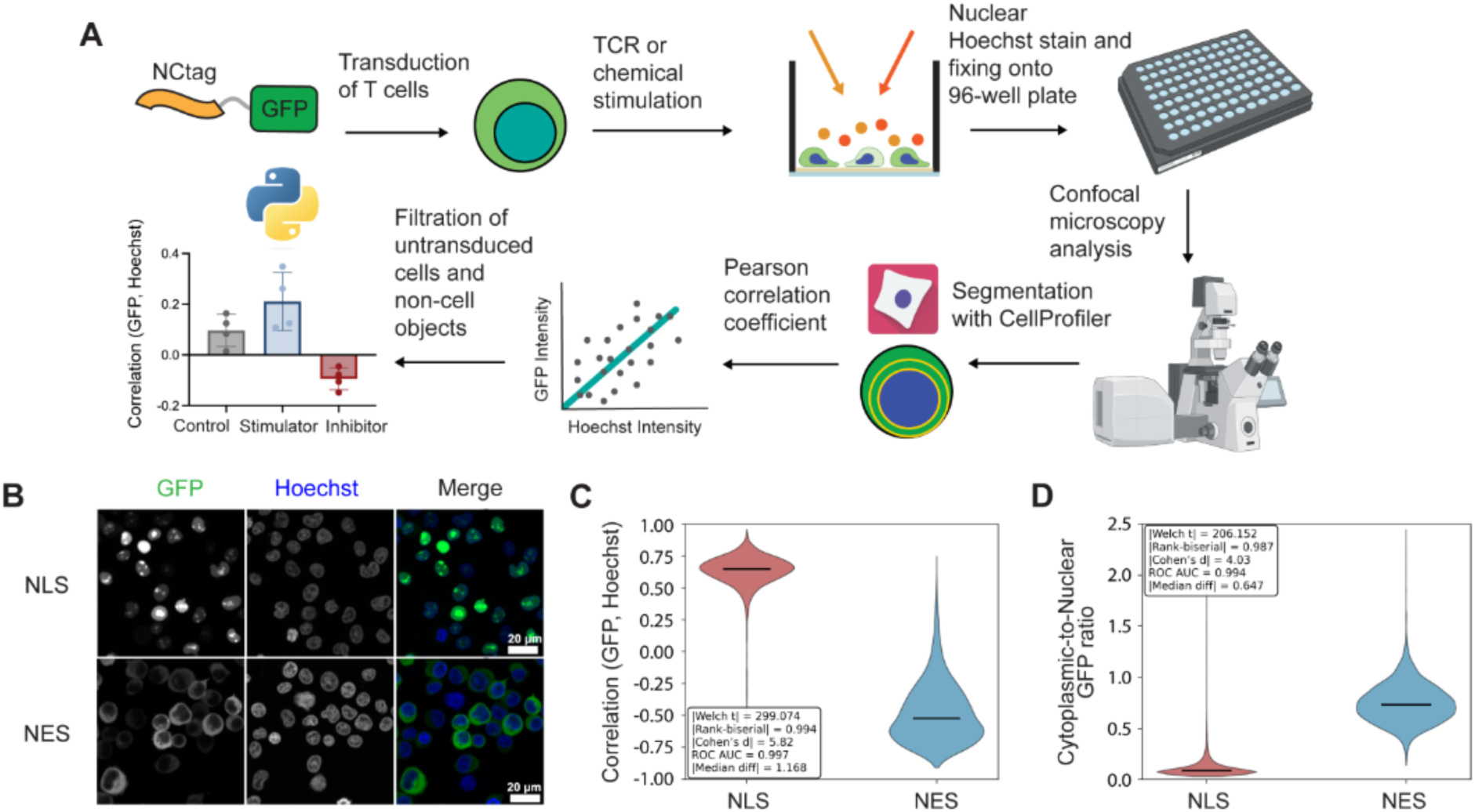
Quantification of NCtag localization in T cells. **A** Schematic of the workflow for assessing the localization of GFP-tagged NCtags in T cells. **B** Representative images of one donor of primary T cells expressing GFP fused to a 3xNLS or 3xNES. **C** Single-cell correlation of GFP and Hoechst and **D** Cytoplasmic-to-nuclear GFP ratio from confocal microscopy of one donor of primary T cells. Violins represent >1,500 cells per condition from one donor; horizontal line indicates median. Insets: statistical analysis.

**Supplementary Figure 2.**
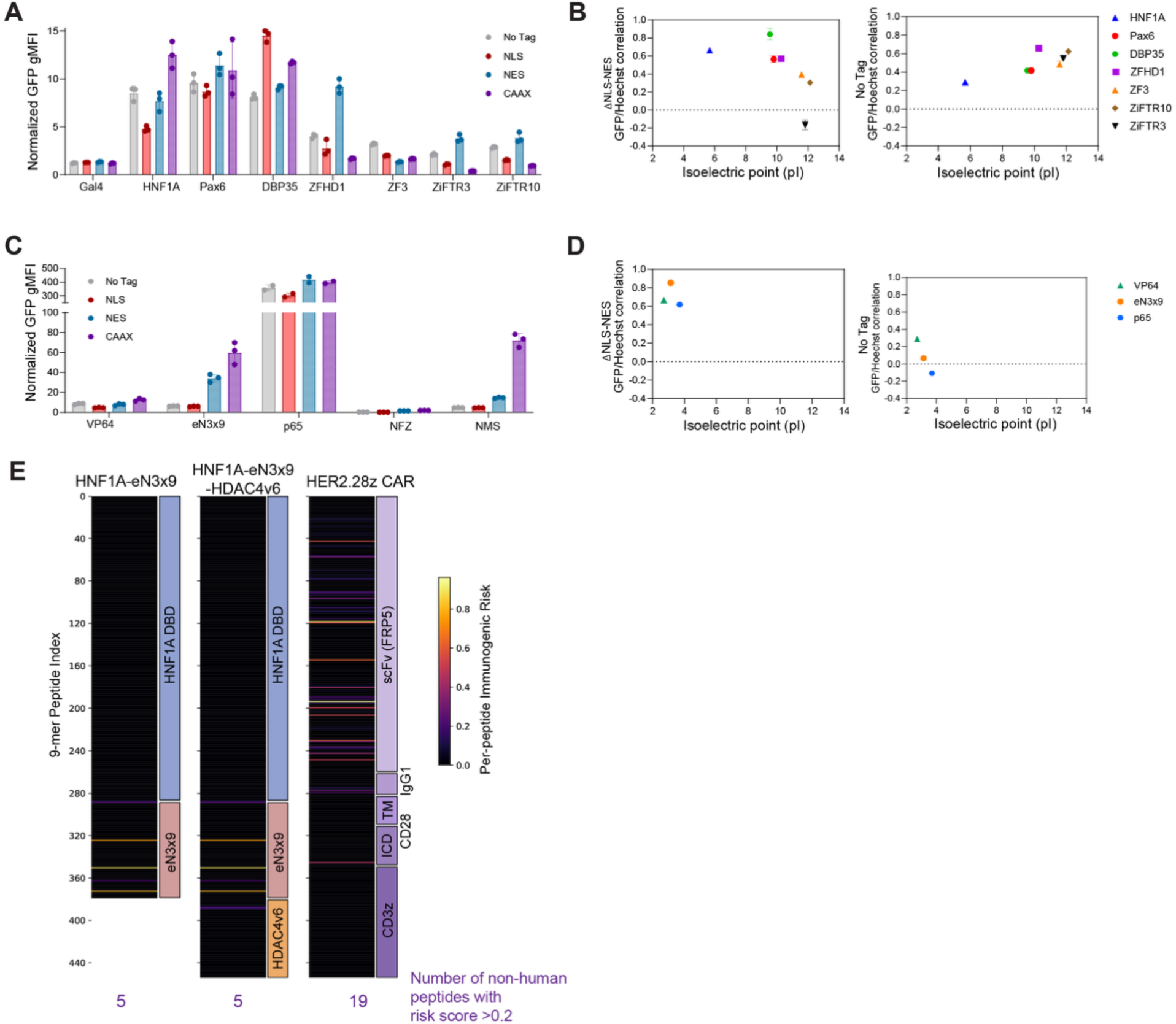
Expression and localization dependencies of synTF variants. **A** Expression of DBDs fused to GFP, VP64, and the indicated localization tags. **B** Difference in average GFP/Hoechst correlation between NLS and NES controls (ΔNLS−NES, left) and No Tag GFP/Hoechst correlation (right) for each DBD, plotted as a function of theoretical isoelectric point (pI) of the DBD. **C** Expression of HNF1A-DBD fused to a panel of TADs, GFP, and the indicated localization tags. (**A,C**) GFP gMFI normalized to untransduced; n = 3 separate transductions in Jurkat cells (2 for p65). **D** Difference in GFP/Hoechst correlation between NLS and NES controls (ΔNLS−NES, left) and No Tag GFP/Hoechst correlation (right) for each TAD, plotted as a function of theoretical pI of the TAD. **E** Immunogenic burden score plotted for each 9-mer peptide based on predicted MHC class I immunogenicity from NETMHCpan4.1 (see Methods for score definition). Human peptides have a score of 0, and the total number of peptides with a risk score >0.2 is given below each protein.

**Supplementary Figure 3.**
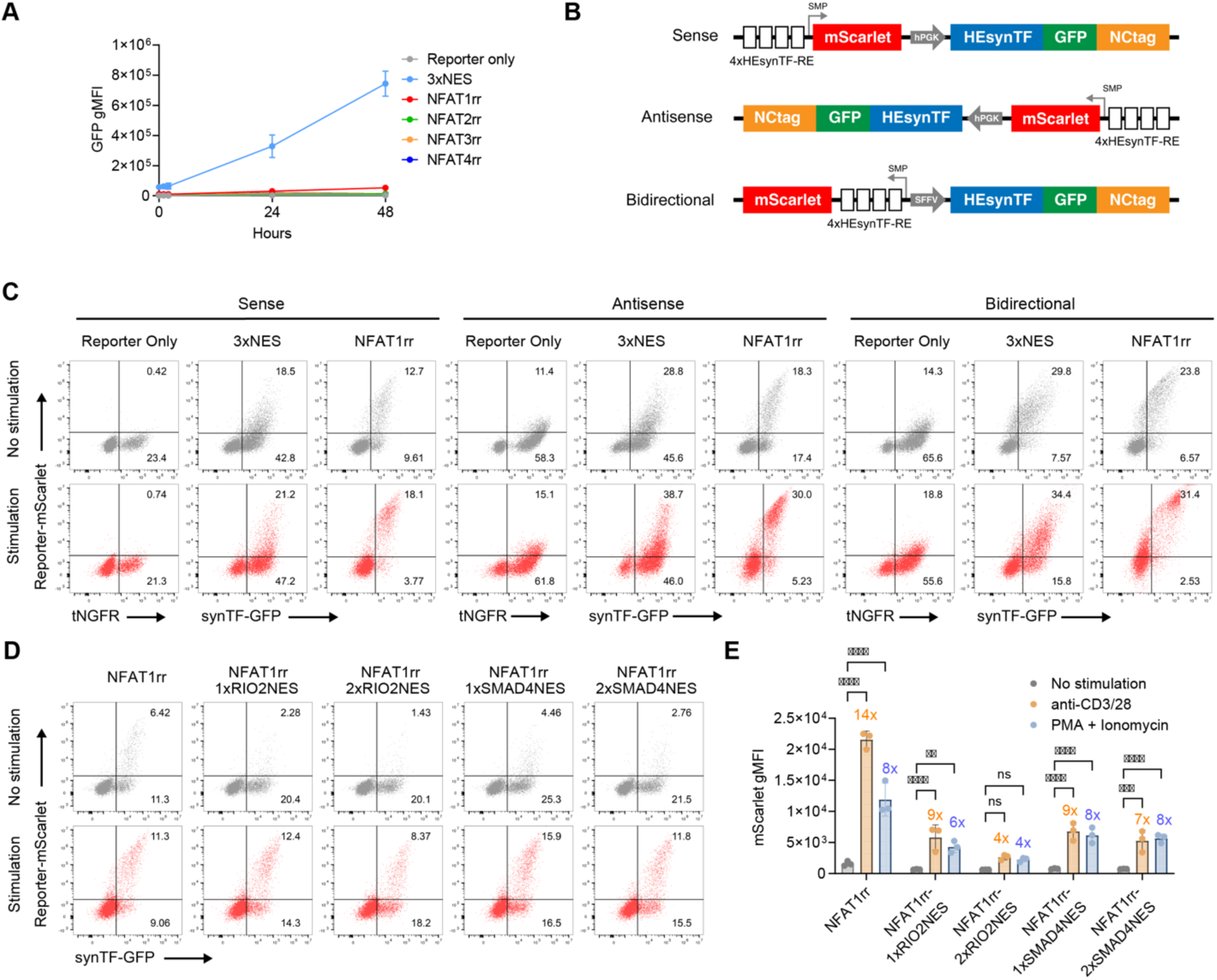
Expression of NFATrr NCtags and engineering to minimize background induction. All panels show primary T cells. **A** Expression of HEsynTF-NFAT1–4rr constructs fused to GFP over a time course of stimulation with anti-CD3 and anti-CD28. Mean ± SD of n = 3 donors. **B** Schematic of the configurations of all-in-one lentiviral vectors tested. **C** Representative flow cytometry dot plots of GFP expression and mScarlet reporter ouput of the indicated constructs with different lentiviral architectures of one donor after 24 h stimulation with anti-CD3/28. **D** Representative flow cytometry dot plots of cells expressing HEsynTF-NFAT1rr fused to different NES sequences of one donor after 24 h stimulation with anti-CD3/CD28. **E** Reporter mScarlet gMFI output of GFP⁺ cells expressing HEsynTF-NFAT1rr fused to different NES sequences after 24 h stimulation with the indicated treatments. n = 3 donors, error bars show SD; significance calculated by two-way ANOVA with Dunett’s adjustment for comparisons vs. no stimulation; **p < 0.01, ***p < 0.001, ****p < 0.0001. (D-E Constructs with WPRE removed from the vector)

**Supplementary Figure 4.**
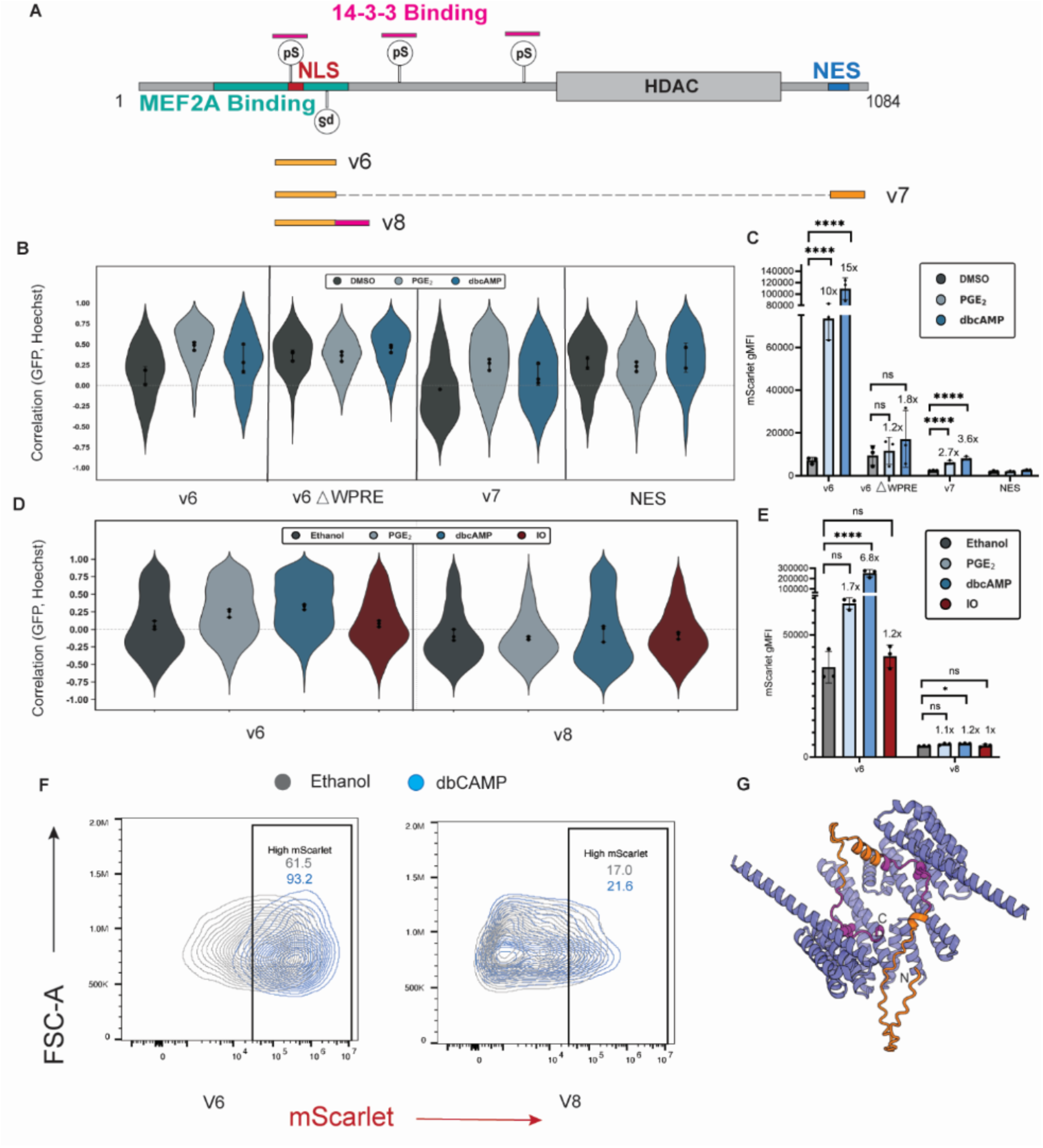
Strategies to reduce basal nuclear localization for HDAC4 NCtag. **A** Schematic representation of the HDAC4 v7 and v8 NCtags. **B** Violin plots quantified from confocal images to measure NCtag localization in v6, v6 ΔWPRE, v7 and 3xNES after the indicated treatments for 24 hours (500 μM dbcAMP or 1 μM PGE_2_) control for >1,360 cells per condition; dots indicate donor means (n = 3 donors); error bars show SD. **C** mScarlet reporter output quantified by flow cytometry for HDAC4 v6 and v7 NCtags fused to HEsynTF after the indicated treatments for 24 hours (500 μM dbcAMP or 1 μM PGE_2_). dots indicate donor means (n = 3 donors); error bars show SD. Significance across treatment conditions was evaluated using a linear mixed-effects model with donor identity specified as a random intercept, followed by pairwise Wald contrasts with Holm-Bonferroni step-down correction (*p < 0.05,**p < 0.01, ***p < 0.001, ****p < 0.0001, ns = not significant). **D** Violin plots quantified from confocal images to measure NCtag localization in v6 and v8 for > 5800 cells per condition for 24-hour treatments (500 μM dbcAMP, 1 μM PGE2, 1.34 μM IO); dots indicate mean of replicates (n = 3 treatment replicates, Jurkat cells); error bars show SD. **E** Reporter output for HEsynTF fused to v6 and v8 NCtags in response to indicated treatments (500 μM dbcAMP, 1 μM PGE_2_, 1.34 μM IO), error bars show SD for all panels. Significance across treatment conditions was evaluated using one way-ANOVA followed by pairwise linear contrasts with Holm-Bonferroni step-down correction for multiple testing (*p < 0.05,**p < 0.01, ***p < 0.001, ****p < 0.0001, ns = not significant). **F** Contour plot of 24-hour vehicle ethanol vs. dbcAMP treatments for HEsynTF fused to v6 (left) and v8 (right), NCtags show extended tail towards mScarlet region for dbCAMP-treated cells. **G** AlphaFold 3 predicted assembly of 14-3-3 dimer (lavender) bound to HDAC4 v8 NCtag (orange), highlighting the primary contact residues (magenta) with the 14-3-3 dimer (pSer residues in spheres).

**Supplementary Figure 5.**
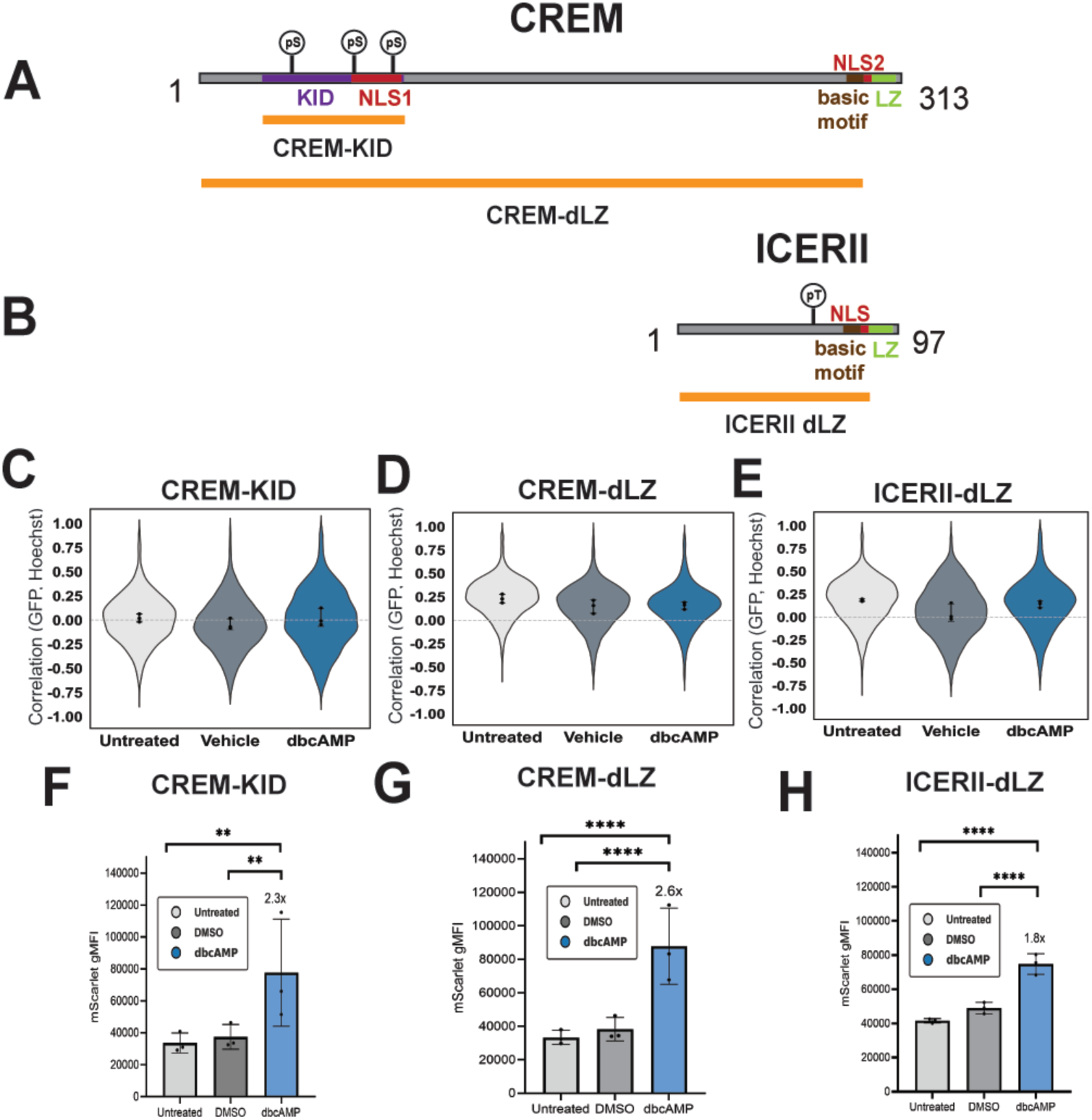
cAMP-responsive NCtags derived from CREM variants. **A** Schematic of full-length CREM, indicating the kinase-inducible domain (KID), NLS, leucine zipper (LZ), and phosphoserines (pS). Orange bars indicate the regions used in the CREM-KID and CREM-dLZ NCtag constructs. **B** Schematic of ICERII, indicating the NLS, LZ, and pS. The orange bar indicates the region used in the ICERII-dLZ NCtag construct. Violin plots of single-cell GFP–Hoechst correlation quantified from confocal microscopy images of Jurkat T cells expressing **C** CREM-KID, **D** CREM-dLZ, and **E** ICERII-dLZ NCtags under untreated, vehicle (DMSO), and 500 μM dbcAMP treatment conditions at 24 hours. Violins represent >3,600 cells per condition; dots indicate per-replicate means (n = 3 independent treatment replicates); error bars show SD. Reporter mScarlet gMFI output in Jurkat T cells expressing HEsynTF fused to **F** CREM-KID, **G** CREM-dLZ, and **H** ICERII-dLZ NCtags under untreated, vehicle (DMSO), and 500 μM dbcAMP conditions at 24 hours. Dots indicate per-replicate means (n = 3 independent treatment replicates); error bars show SD. Significance across treatment conditions was evaluated using a linear mixed-effects model with experimental day specified as a random intercept, followed by pairwise Wald contrasts with Holm-Bonferroni step-down correction (*p < 0.05,**p < 0.01, ***p < 0.001, ****p < 0.0001, ns = not significant).

**Supplementary Figure 6.**
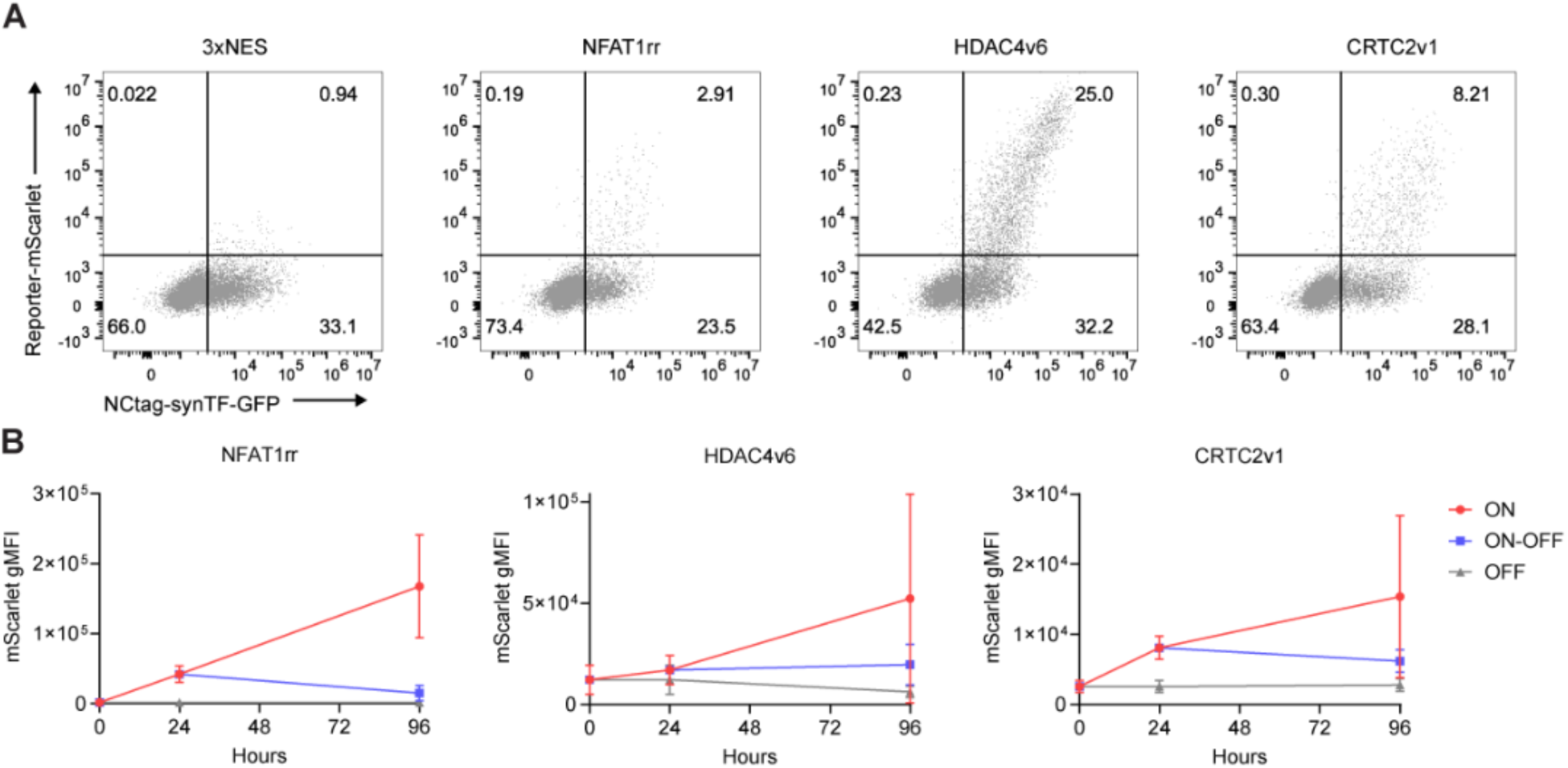
Temporal control of NCtag-synTF reporter output with sustained, absent, and transient inputs. All panels show primary T cells expressing HEsynTF fused to the indicated NCtags. **A** Representative flow cytometry dot plots depicting GFP expression and mScarlet reporter output of one donor at baseline (OFF) levels. **B** Reporter mScarlet gMFI output of cells stimulated or unstimulated for the indicated time. NFAT1rr was stimulated with plate-bound anti-CD3 and anti-CD28; HDAC4v6 and CRTC2v1 were stimulated with 500 μM dbcAMP. Mean ± SD of n = 4 donors.

**Supplementary Figure 7.**
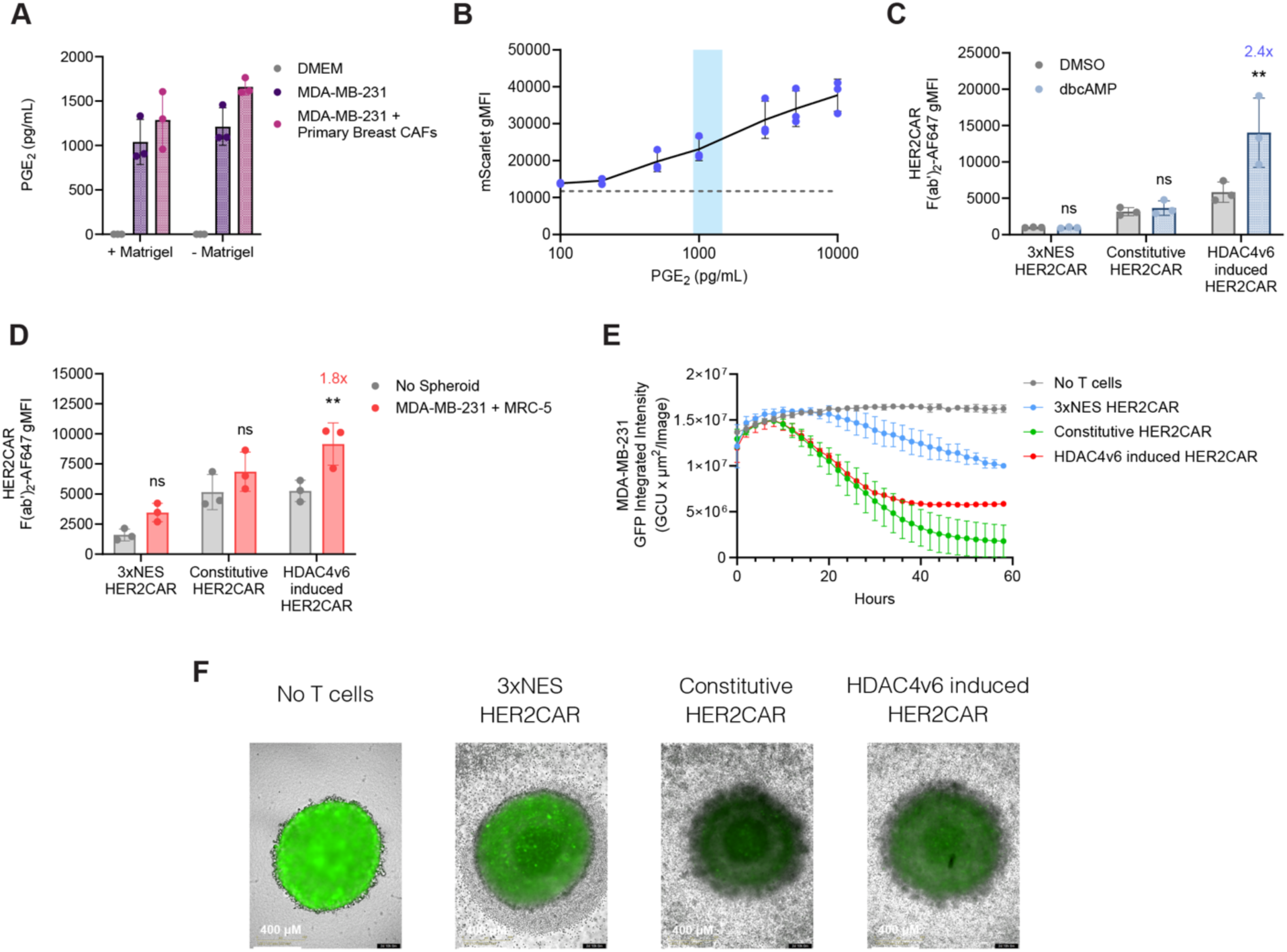
Application of HDAC4v6 NCtag to control HER2CAR expression with endogenous PGE_2_ in vitro. All panels show primary T cells, unless otherwise noted. **A** ELISA quantification of PGE_2_ in MDA-MB-231 spheroids with or without primary cancer-associated fibroblasts and Matrigel. n = 3 biological replicates; error bars show SD. **B** Reporter mScarlet gMFI output of HEsynTF-HDAC4v6 in GFP⁺ cells under a PGE_2_ titration, quantified at 24 hr. n = 3 donors; error bars show SD. Blue shaded region corresponds to PGE_2_ levels measured in MDA-MB-231 spheroids in (A) and dotted line to baseline levels of mScarlet gMFI. **C** HER2 CAR expression (gMFI) in GFP⁺ cells induced by HEsynTF-3xNES, HEsynTF-HDAC4v6, or a constitutive EFS promoter after 24h treatment with 500 µM db-cAMP. **D** HER2 CAR expression (F(ab’)_2_ - AlexaFluor647 gMFI) of GFP⁺ cells induced by HEsynTF-3xNES, HEsynTF-HDAC4v6, or a constitutive EFS promoter after 58h co-culture with MDA-MB-231 and MRC-5 tumor spheroids. C-D n = 3 donors; error bars show SD; significance calculated by two-way ANOVA with Bonferroni’s adjustment for multiple comparisons; **p< 0.01. **E** MDA-MB-231-GFP integrated intensity over time during co-culture with primary T cells expressing the indicated constructs. Mean ± SD of n = 3 donors. **F** Representative images of MDA-MB-231-GFP spheroids after 58h co-culture with T cells expressing HER2 CAR under the indicated constructs.

## References

1. Bhatt, B., García-Díaz, P. & Foight, G. W. Synthetic transcription factor engineering for cell and gene therapy. Trends Biotechnol. 42, 449–463 (2024).

2. Lee, S., Khalil, A. S. & Wong, W. W. Recent progress of gene circuit designs in immune cell therapies. Cell Syst. 13, 864–873 (2022).

3. Galvan, S., Madderson, O., Xue, S., Teixeira, A. P. & Fussenegger, M. Regulation of Transgene Expression by the Natural Sweetener Xylose. Adv. Sci. 9, 1–10 (2022).

4. Shao, J. et al. Smartphone-controlled optogenetically engineered cells enable semiautomatic glucose homeostasis in diabetic mice. Sci. Transl. Med. 9, eaal2298 (2017).

5. Bojar, D., Scheller, L., Hamri, G. C. E., Xie, M. & Fussenegger, M. Caffeine-inducible gene switches controlling experimental diabetes. Nat. Commun. 9, (2018).

6. Mansouri, M. et al. Smart-watch-programmed green-light-operated percutaneous control of therapeutic transgenes. Nat. Commun. 12, (2021).

7. Barrett, J. et al. A Synthetic Biology Rheoswitch Therapeutic System® for the Controlled Local Expression of IL-12 as an Immunotherapy for the Treatment of Cancer. Cell Biol. Res. Ther. 5, (2016).

8. Manhas, J., Edelstein, H. I., Leonard, J. N. & Morsut, L. The evolution of synthetic receptor systems. Nat. Chem. Biol. 18, 244–255 (2022).

9. Roybal, K. T. et al. Precision Tumor Recognition by T Cells With Combinatorial Antigen-Sensing Circuits. Cell 164, 770–779 (2016).

10. Zhu, I. et al. Modular design of synthetic receptors for programmed gene regulation in cell therapies. Cell 185, 1431–1443.e16 (2022).

11. Allen, G. M. et al. Synthetic cytokine circuits that drive T cells into immune-excluded tumors. Science 378, eaba1624 (2022).

12. Piraner, D. I. et al. Engineered receptors for soluble cellular communication and disease sensing. Nature 638, 805–813 (2025).

13. Krawczyk, K., Scheller, L., Kim, H. & Fussenegger, M. Rewiring of endogenous signaling pathways to genomic targets for therapeutic cell reprogramming. Nat. Commun. 11, 1–9 (2020).

14. Campbell, B. C. et al. mGreenLantern: a bright monomeric fluorescent protein with rapid expression and cell filling properties for neuronal imaging. Proc. Natl. Acad. Sci. 117, 30710–30721 (2020).

15. Hwang, J. R., Byeon, Y., Kim, D. & Park, S. G. Recent insights of T cell receptor-mediated signaling pathways for T cell activation and development. Exp. Mol. Med. 52, 750–761 (2020).

16. Gross, S. M. & Rotwein, P. Akt signaling dynamics in individual cells. J. Cell Sci. 128, 2509–2519 (2015).

17. Meško, M., Lebar, T., Dekleva, P., Jerala, R. & Benčina, M. Engineering and Rewiring of a Calcium-Dependent Signaling Pathway. ACS Synth. Biol. 9, 2055–2065 (2020).

18. Regot, S., Hughey, J. J., Bajar, B. T., Carrasco, S. & Covert, M. W. High-sensitivity measurements of multiple kinase activities in live single cells. Cell 157, 1724–1734 (2014).

19. Guo, T., Ma, D. & Lu, T. K. Sense-and-Respond Payload Delivery Using a Novel Antigen-Inducible Promoter Improves Suboptimal CAR-T Activation. ACS Synth. Biol. 11, 1440–1453 (2022).

20. Merlet, E. et al. A calcium-sensitive promoter construct for gene therapy. Gene Ther. 20, 248–254 (2013).

21. Roscilli, G. et al. Long-term and tight control of gene expression in mouse skeletal muscle by a new hybrid human transcription factor. Mol. Ther. 6, 653–663 (2002).

22. Glasscock, C. J. et al. Computational design of sequence-specific DNA-binding proteins. Nat. Struct. Mol. Biol. 32, 2252–2261 (2025).

23. Donahue, P. S. et al. The COMET toolkit for composing customizable genetic programs in mammalian cells. Nat. Commun. 11, (2020).

24. Pomerantz, J. L., Sharp, P. A. & Pabo, C. Structure-Based Design of Transcription Factors. Science 267, 93–96 (1995).

25. Li, H. S. et al. Multidimensional control of therapeutic human cell function with synthetic gene circuits. Science 378, 1227–1234 (2022).

26. Mahata, B. et al. Compact engineered human mechanosensitive transactivation modules enable potent and versatile synthetic transcriptional control. Nat. Methods 20, 1716–1728 (2023).

27. Tycko, J. et al. Development of compact transcriptional effectors using high-throughput measurements in diverse contexts. Nat. Biotechnol. 43, 1525–1538 (2025).

28. Marchingo, J. M. & Cantrell, D. A. Protein synthesis, degradation, and energy metabolism in T cell immunity. Cell. Mol. Immunol. 19, 303–315 (2022).

29. Fu, S.-C., Fung, H. Y. J., Cağatay, T., Baumhardt, J. & Chook, Y. M. Correlation of CRM1-NES affinity with nuclear export activity. Mol. Biol. Cell 29, 2037–2044 (2018).

30. Lee, S. D., Colla, E., Sheen, M. R., Na, K. Y. & Kwon, H. M. Multiple Domains of TonEBP Cooperate to Stimulate Transcription in Response to Hypertonicity. J. Biol. Chem. 278, 47571–47577 (2003).

31. Khandwala, C. B. et al. Direct ionic stress sensing and mitigation by the transcription factor NFAT5. Sci. Adv. 11, (2025).

32. López-Rodríguez, C. et al. Bridging the NFAT and NF-κB families: NFAT5 dimerization regulates cytokine gene transcription in response to osmotic stress. Immunity 15, 47–58 (2001).

33. Lacher, S. B. et al. PGE2 limits effector expansion of tumour-infiltrating stem-like CD8+ T cells. Nature 629, 417–425 (2024).

34. Morotti, M. et al. PGE2 inhibits TIL expansion by disrupting IL-2 signalling and mitochondrial function. Nature 629, 426–434 (2024).

35. Globig, A.-M. et al. The β1-adrenergic receptor links sympathetic nerves to T cell exhaustion. Nature 622, 383–392 (2023).

36. Huang, S., Apasov, S., Koshiba, M. & Sitkovsky, M. Role of A2a extracellular adenosine receptor-mediated signaling in adenosine-mediated inhibition of T-cell activation and expansion. Blood 90, 1600–1610 (1997).

37. Helmstadter, K. G. et al. CaMKII and PKA-dependent phosphorylation co-regulate nuclear localization of HDAC4 in adult cardiomyocytes. Basic Res. Cardiol. 116, 1–17 (2021).

38. Screaton, R. A. et al. The CREB Coactivator TORC2 Functions as a Calcium- and cAMP-Sensitive Coincidence Detector. Cell 119, 61–74 (2004).

39. Vaeth, M. et al. Regulatory T cells facilitate the nuclear accumulation of inducible cAMP early repressor (ICER) and suppress nuclear factor of activated T cell c1 (NFATc1). Proc. Natl. Acad. Sci. 108, 2480–2485 (2011).

40. Liu, Y. & Schneider, M. F. Opposing HDAC4 nuclear fluxes due to phosphorylation by β-adrenergic activated protein kinase A or by activity or Epac activated CaMKII in skeletal muscle fibres. J. Physiol. 591, 3605–3623 (2013).

41. Wang, A. H. & Yang, X.-J. Histone Deacetylase 4 Possesses Intrinsic Nuclear Import and Export Signals. Mol. Cell. Biol. 21, 5992–6005 (2001).

42. Zhang, H., Kong, Q., Wang, J., Jiang, Y. & Hua, H. Complex roles of cAMP–PKA–CREB signaling in cancer. Exp. Hematol. Oncol. 9, 1–13 (2020).

43. Chow, C.-W. & Davis, R. J. Integration of Calcium and Cyclic AMP Signaling Pathways by 14-3-3. Mol. Cell. Biol. 20, 702–712 (2000).

44. Schrey, M. P. & Patel, K. V. Prostaglandin E2 production and metabolism in human breast cancer cells and breast fibroblasts. Regulation by inflammatory mediators. Br. J. Cancer 72, 1412–9 (1995).

45. Kock, A. et al. Inhibition of Microsomal Prostaglandin E Synthase-1 in Cancer-Associated Fibroblasts Suppresses Neuroblastoma Tumor Growth. EBioMedicine 32, 84–92 (2018).

46. Caronni, N. et al. IL-1β+ macrophages fuel pathogenic inflammation in pancreatic cancer. Nature 623, 415–422 (2023).

47. Rafei, H. et al. CREM is a regulatory checkpoint of CAR and IL-15 signalling in NK cells. Nature 643, 1076–1086 (2025).

48. Tillé, L. et al. Activation of the transcription factor NFAT5 in the tumor microenvironment enforces CD8+ T cell exhaustion. Nat. Immunol. 24, 1645–1653 (2023).

49. Leslie, T. K. et al. Sodium homeostasis in the tumour microenvironment. Biochim. Biophys. Acta - Rev. Cancer 1872, (2019).

50. Kosti, P. et al. Hypoxia-sensing CAR T cells provide safety and efficacy in treating solid tumors. Cell Rep. Med. 2, (2021).

51. Manchon, E. et al. Harnessing nutrient scarcity for enhanced CAR-T-cell potency and safety in solid tumors. Cell. Mol. Immunol. 22, 645–660 (2025).

52. Halterman, J. A., Moo Kwon, H. & Wamhoff, B. R. Tonicity-independent regulation of the osmosensitive transcription factor TonEBP (NFAT5). Am J Physiol Cell Physiol 302, 1–8 (2012).

53. Kim, N., Shin, S. & Bae, S. W. cAMP Biosensors Based on Genetically Encoded Fluorescent/Luminescent Proteins. Biosensors 11, 39 (2021).

54. Kudo, T. et al. Live-cell measurements of kinase activity in single cells using translocation reporters. Nat. Protoc. 13, 155–169 (2018).

55. Ahmed, N. et al. Immunotherapy for osteosarcoma: Genetic modification of T cells overcomes low levels of tumor antigen expression. Mol. Ther. 17, 1779–1787 (2009).

56. Kochenderfer, J. et al. Construction and Pre-Clinical Evaluation of An Anti-CD19 Chimeric Antigen Receptor. Blood 112, 4623–4623 (2008).

57. Watanabe, N. & McKenna, M. K. Generation of CAR T-Cells Using γ-Retroviral Vector. Methods in Cell Biology vol. 167 (Elsevier Inc., 2022).

58. Han, J., Tam, K., Tam, C., Hollis, R. P. & Kohn, D. B. Improved lentiviral vector titers from a multi-gene knockout packaging line. Mol. Ther. - Oncolytics 23, 582–592 (2021).

59. Zhang, X.-Y., La Russa, V. F. & Reiser, J. Transduction of Bone-Marrow-Derived Mesenchymal Stem Cells by Using Lentivirus Vectors Pseudotyped with Modified RD114 Envelope Glycoproteins. J. Virol. 78, 1219–1229 (2004).

60. Bäckström, A. et al. A Sample Preparation Protocol for High Throughput Immunofluorescence of Suspension Cells on an Adherent Surface. J. Histochem. Cytochem. 68, 473–489 (2020).

61. Stirling, D. R. et al. CellProfiler 4: improvements in speed, utility and usability. BMC Bioinformatics 22, (2021).

62. Schneider, C. A., Rasband, W. S. & Eliceiri, K. W. NIH Image to ImageJ: 25 years of image analysis. Nat. Methods 9, 671–675 (2012).

63. Reynisson, B., Alvarez, B., Paul, S., Peters, B. & Nielsen, M. NetMHCpan-4.1 and NetMHCIIpan-4.0: improved predictions of MHC antigen presentation by concurrent motif deconvolution and integration of MS MHC eluted ligand data. Nucleic Acids Res. 48, W449–W454 (2020).

